# JunB-HBZ nuclear translocation by TGF-β is a key driver in HTLV-1–mediated leukemogenesis

**DOI:** 10.1101/2024.10.06.616923

**Authors:** Wenyi Zhang, Takafumi Shichijo, Xueda Chen, Masao Matsuoka, Jun-ichirou Yasunaga

**Author notes:** Correspondence to Takafumi Shichijo, Jun-ichirou Yasunaga **Email:**.

## Abstract

The *HTLV-1 bZIP factor* (*HBZ*) gene, which is the only viral gene conserved and consistently expressed in all adult T-cell leukemia-lymphoma (ATL) cases, is critical for ATL oncogenesis. Although HBZ protein is found in both the nucleus and the cytoplasm, the dynamics of HBZ protein localization and its contribution to oncogenesis have not been fully elucidated. In this study, we analyzed the subcellular expression pattern of HBZ in primary HTLV-1–infected T cells from asymptomatic carriers and leukemic cells of ATL patients using the Proximity Ligation Assay. Nuclear localization of HBZ protein was significantly higher in fresh ATL cells than in HTLV-1–infected cells from carriers. Importantly, translocation of HBZ protein from the cytoplasm to the nucleus after TGF-β activation was observed in ATL patients, but not in HTLV-1 carriers. In ATL cells, the cellular transcription factors JunB and pSmad3 interact with HBZ and facilitate its nuclear translocation upon TGF-β stimulation. *JUNB* knockdown inhibits cell proliferation *in vitro* and *in vivo* and promotes apoptosis in ATL cells but not in HTLV-1–infected non-leukemic cells, indicating that JunB has important roles in maintaining ATL cells. In conclusion, TGF-β-induced nuclear translocation of HBZ-JunB complexes is associated with ATL oncogenesis.

## Introduction

The signaling induced by transforming growth factor β (TGF-β) has pleiotropic actions in a wide range of cellular processes, such as cell differentiation, proliferation, apoptosis, secretion of extracellular mediators, tissue fibrosis and epithelial-to-mesenchymal transition (EMT). When the TGF-β signaling pathway is activated, Smad3 protein is phosphorylated and forms a complex with Smad4, which then translocates to the nucleus to regulate the transcriptional activation of various genes^1^. The functions of TGF-β are dependent not only on cell type or cancer type but also on the particular stage of tumorigenesis. In the immune system, TGF-β exerts inhibitory effects in multiple ways^2^. TGF-β inhibits Th1 and Th2 differentiation, suppresses the trafficking and cytotoxic activities of CD8+ T cells and induces the differentiation of regulatory T cells (Tregs). Thus, activating the TGF-β signaling pathway enables cancer cells to evade host immunity. Notably, however, little is known about the role of TGF-β in T-cell malignancies.

Human T-cell leukemia virus type 1 (HTLV-1) is the first discovered human oncogenic retrovirus that primarily infects CD4^+^ T cells^3, 4^. While most HTLV-1– infected individuals remain asymptomatic for life, HTLV-1 does induce a malignancy of CD4^+^ T cells, named adult T-cell leukemia-lymphoma (ATL), in 2–5% of carriers after several decades of latency^5, 6^. Viral replication of HTLV-1 is typically suppressed *in vivo*, and persistent infection is established primarily through clonal proliferation of infected cells^7^.

Among the HTLV-1 viral proteins, HTLV-1 basic leucine zipper factor (HBZ), which is encoded in the antisense strand of the provirus, is consistently expressed in all ATL cases^8, 9^. Knockdown of *HBZ* leads to suppression of ATL cell proliferation^7^ and significantly reduces tumor formation and tissue infiltration by xenografts transplanted into immunocompetent mice^8, 10^, indicating the potential necessity of *HBZ* in the oncogenesis of ATL. It is noteworthy that HBZ increases the number of induced Treg (iTreg) cells with high Foxp3 expression^11^ by activating the TGF-β/Smad signaling pathway^12^, which plays crucial roles not only in immunosuppression but also in promoting the proliferation of ATL cells^13^. Thus HBZ-mediated activation of the TGF-β/Smad signaling pathway has a special impact on ATL leukemogenesis.

The HBZ protein has been reported to be localized in the nucleus when it is transiently overexpressed^12, 14^. We previously reported that subcellular localization of HBZ protein is changed by cell context, and HBZ protein can be localized in both nucleus and cytoplasm in T cells^15^. Indeed, it has been reported that HBZ protein tends to be present in both the cytoplasm and the nucleus in ATL cells, while it is expressed in the cytoplasm of infected cells from HTLV-1 carriers^16, 17^. HBZ contains three nuclear localization signals and a functional nuclear export signal sequence in its N-terminal region, indicating that it can be shuttled between the cytoplasm and the nucleus^18, 19^. HBZ interacts with various cellular factors, controlling their activities in both the nucleus and cytoplasm^15, 19^. However, it remains unclear how the subcellular localization of HBZ is regulated and what significance this may have in the oncogenesis of ATL.

In the present study, to evaluate intracellular localization of endogenous HBZ protein in fresh ATL/HTLV-1–infected cells, we employ a more precise and highly sensitive method, the proximity ligation assay (PLA). The results demonstrate that HBZ protein in fresh ATL cells exhibits a higher nuclear localization than in infected cells from carriers. Surprisingly, in response to TGF-β stimulation, HBZ protein translocates from the cytoplasm to the nucleus in ATL cells, but not in non-leukemic infected cells. Importantly, an AP-1 protein, JunB, which is a known binding partner of HBZ, was indispensable for its translocation to the nucleus and the exertion of the cell proliferation-promoting functions triggered by TGF-β stimulation. Notably, we also found that JunB is essential for ATL tumorigenesis *in vivo*. These results suggest that the link between the TGF-β/Smad pathway and translocation of the HBZ-JunB complex plays a master regulatory role in ATL oncogenesis.

## Results

### A higher fraction of HBZ is nuclear in ATL cells than in HTLV-1–infected cells

We first analyzed the expression patterns of HBZ in ATL cell lines (MT1, ATL43T[+], ED, and TL-Om1), HTLV-1 immortalized cell lines (MT-4, and Hut102), and HTLV-1–infected cell lines derived from non-leukemic clones (ATL6, and ATL7)^20^ using the Duolink^®^ PLA. Fluorescence signals were objectively and quantitatively evaluated by ImageJ, and the proportion of nuclear signals in the total signal was calculated for each cell. Unlike transiently overexpressed HBZ, which is exclusively found in the nucleus^12, 15^, HBZ in all the cell lines was distributed in both the nucleus and the cytoplasm, though the nuclear-cytoplasmic ratio varied (**Fig. 1a, b**, and Supplementary Fig. 1a). Compared to the ATL cell lines, HBZ exhibited a stronger tendency for cytoplasmic expression in the two HTLV-1–infected non-leukemic cell lines, indicating that the intracellular localization of HBZ differs between leukemic and non-leukemic cells. Notably, in the MT-1 cell line, approximately 60% of the HBZ protein signals are distributed in the nucleus. MT-1 has an equivalent viral gene expression profile to primary HTLV-1–infected cells^21, 22^. This led us to wonder whether the expression of HBZ in fresh ATL patient samples and HTLV-1–infected carrier samples might also differ in a similar way.

**Fig. 1.**
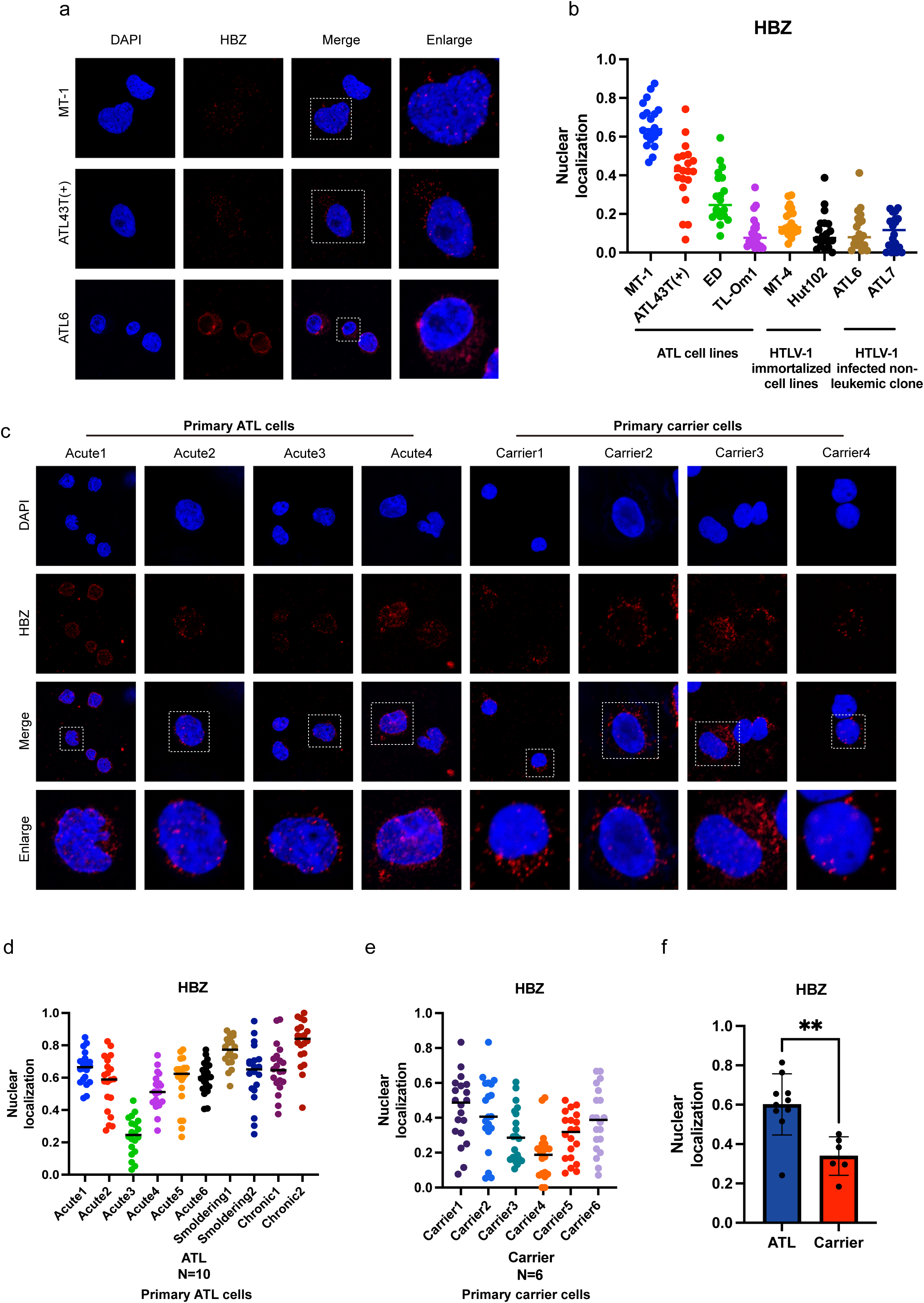
Intracellular localization pattern of HBZ protein in various cell lines and patient samples. **a** Localization of HBZ protein in MT-1, ATL43T (+) and ATL6 cell lines by Duolink^®^ PLA. **b** The proportion of HBZ that is localized to the nucleus (i.e. nuclear HBZ signals divided by total HBZ signals) is shown for ATL cell lines, HTLV-1 immortalized cell lines and HTLV-1–infected cell lines derived from non-leukemic clones, which were analyzed by ImageJ. 20 randomly selected cells were analyzed for each cell line. **c** Localization of HBZ protein in primary cells from ATL patients (n = 4) and HTLV-1– infected carriers (n = 4) by Duolink^®^ PLA. **d, e** The proportion of HBZ that is localized to the nucleus in each ATL patient sample (n = 10) and HTLV-1–infected carrier sample (n = 6) was analyzed by ImageJ. 20 randomly selected cells were analyzed for each patient. **f** The proportion of HBZ that is localized to the nucleus was compared between ATL (N = 10) and HTLV-1–infected carrier (N = 6) samples (two-tailed unpaired Student’s t test; **P < 0.01). All experiments were performed at least twice.

Therefore we examined HBZ expression in patient samples, including ten ATL patient samples (six acute-type, two smoldering-type, and two chronic-type), and six HTLV-1–infected carrier samples. In the ATL cases, the distribution of HBZ protein was throughout the cytoplasm and nucleus and the localization was usually higher in the nucleus (**Fig. 1c, left, Fig. 1d**, and Supplementary Fig. 1b, c). In contrast, in HTLV-1 carrier cells that expressed HBZ, more HBZ was found in the cytoplasm than in the nucleus (**Fig. 1c, right, Fig. 1e**, and Supplementary Fig. 1b, c). The proportion of nuclear localization of HBZ protein is significantly higher in fresh ATL cells compared to that of HTLV-1–infected cells from HTLV-1 carriers (**Fig. 1f**). Collectively, these data show that the nuclear localization of HBZ protein is associated with the disease status.

### Activation of TGF-β induces translocation of HBZ protein from the cytoplasm to the nucleus

In our previous studies, we demonstrated that signaling via the TGF-β/Smad pathway is enhanced by HBZ^12^, a step which is critical for the oncogenesis of ATL^13^. Activation of TGF-β signaling induces the nuclear localization of phosphorylated Smad3 (pSmad3) protein^13, 23^, which is a known binding partner of HBZ^12^, suggesting that the activation of TGF-β may also change subcellular localization of HBZ.

To investigate whether TGF-β/Smad signaling indeed induces the nuclear translocation of HBZ, we stimulated ATL and HTLV-1–infected non-leukemic cell lines with recombinant TGF-β and observed the sub-localization of HBZ by Duolink^®^ PLA. We found that in (leukemic) TL-Om1 cells, the nuclear proportion of HBZ protein was significantly increased after TGF-β stimulation (**Fig. 2a, b**). To confirm the accumulation of HBZ protein in the nucleus in response to TGF-β stimulation, we performed an immunoblot experiment, separating TL-Om1 cells into cytoplasmic and nuclear fractions. As shown in Supplementary Fig. 2a, the expression of HBZ protein was increased in the nucleus after TGF-β treatment. Furthermore, this shift in HBZ protein expression from the cytoplasm toward the nucleus was also observed in three other ATL cell lines (ED, Hut102 and ATL43T [+]) (**Fig. 2b** and Supplementary Fig. 2b). We next analyzed whether translocation of HBZ can also be induced by TGF-β in two non-leukemic HTLV-1–infected T-cell lines, ATL6 and ATL7. In clear contrast to its effect on ATL cell lines, TGF-β stimulation did not change the intracellular localization of the HBZ protein in either the ATL6 or ATL7 cell line; HBZ remained in the cytoplasm (**Fig. 2c, d** and Supplementary Fig. 2c).

**Fig. 2.**
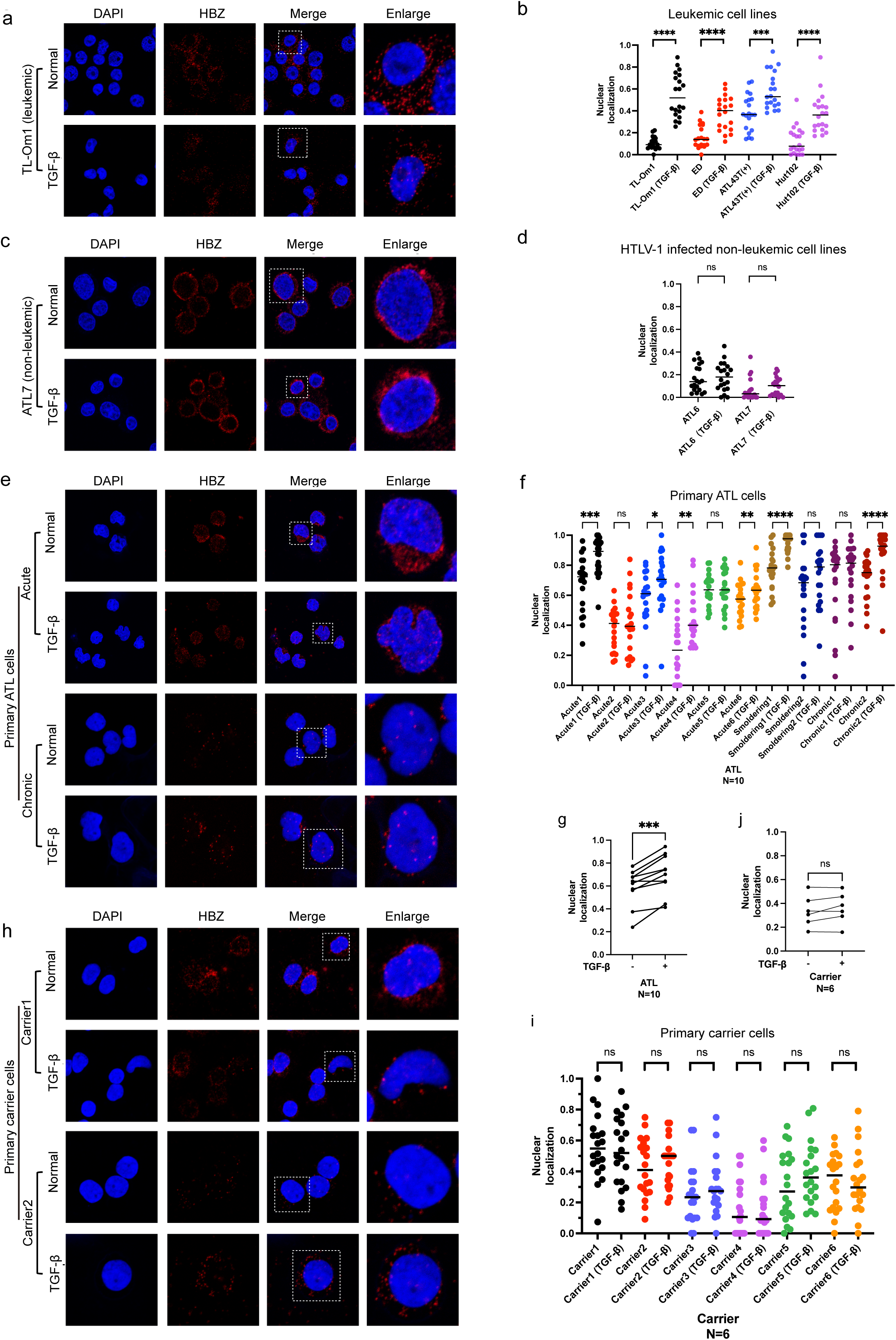
TGF-β treatment induces nuclear translocation of HBZ protein in ATL cell lines and ATL patient samples. **a, c** Localization of HBZ protein in TL-Om1(**a**) and ATL7(**c**) cells with or without TGF-β treatment was detected by Duolink^®^ PLA. **b, d** The proportion of HBZ protein that is localized to the nucleus is shown for ATL cell lines and HTLV-1 immortalized cell lines (**b**) and HTLV-1–infected non-leukemic cell lines (ATL6 and ATL7) (**d**) with or without TGF-β treatment, analyzed using ImageJ. 20 randomly selected cells were analyzed for each cell line. **e** Localization of HBZ protein in primary ATL cells of ATL patients with or without TGF-β treatment. **f-g** The proportion of HBZ protein that is localized to the nucleus in primary ATL cells from 10 ATL patients with or without TGF-β treatment was analyzed individually (**f**) and collectively (**g**) using ImageJ. 20 randomly selected cells were analyzed for each sample. **h** Localization of HBZ protein from primary HTLV-1–infected cells of HTLV-1–infected carrier patients with or without TGF-β treatment. **i, j** The proportion of HBZ protein that is localized to the nucleus in primary HTLV-1–infected cells from 6 HTLV-1–infected carrier patients with or without TGF-β treatment was analyzed individually (**i**) and collectively (**j**) using ImageJ. 20 randomly selected cells were analyzed for each sample. Statistical analyses were performed by two-tailed unpaired Student’s t test. * p < 0.05, ** p < 0.01, *** p < 0.001, **** p < 0.0001, ns. P>0.05. All experiments were performed at least twice.

Next we examined whether TGF-β-induced nuclear transportation of HBZ protein also occurred in fresh ATL or HTLV-1–infected cells obtained from patients or carriers respectively. In the majority of ATL patient samples, HBZ protein was translocated from the cytoplasm to the nucleus in response to TGF-β stimulation (**Fig. 2e-g**). In contrast, all HTLV-1 carrier samples showed no significant change in HBZ localization (**Fig. 2h-j**).

To investigate whether the altered localization of HBZ induced by TGF-β is due to the influence of binding partners, we evaluated the binding of HBZ to pSmad3 by Duolink^®^ PLA. The PLA signal is only detectable when pSmad3 and HBZ proteins interact with each other. As shown in Supplementary Fig. 3a-d, after activation of TGF-β, the HBZ-pSmad3 complex was observed predominantly in the nucleus in the ATL cell line TL-Om1 and in fresh ATL patient samples, whereas it remained in the cytoplasm in the non-leukemic ATL7 cell line. These results suggest that the nuclear localization of HBZ induced by TGF-β in ATL cells may occur via HBZ-Smad3 complexes being transported into the nucleus.

Collectively, our data indicate that HBZ protein shifts its localization from the cytoplasm to the nucleus in response to TGF-β stimulation in ATL cell lines and fresh ATL cells. However, this change in localization is not significantly observed in cell lines derived from HTLV-1–infected non-leukemic clones or in the HTLV-1–infected cells of carriers. These results indicate that the nuclear localization of HBZ protein induced by TGF-β is associated specifically with ATL.

### Increased JunB facilitates the nuclear localization of HBZ in response to TGF-β stimulation

To gain insight into the regulation of TGF-β stimulation in ATL cells, we performed RNA-seq analysis on the ED cell line with or without TGF-β stimulation. Overall, 675 and 656 transcripts were differentially up- or down-regulated in ED cells after 2 h of TGF-β stimulation compared to unstimulated cells (**Fig. 3a**). Consistent with our previous studies^13^, gene set enrichment analysis (GSEA) revealed that the TGF-β signaling pathway affects a range of cell growth-related signaling pathways such as the P53, E2F, apoptosis and G2M pathways, suggesting that the TGF-β signaling pathway is important for ATL cell proliferation. Notably, the MYC-associated gene sets were the most upregulated upon TGF-β stimulation (**Fig. 3b, c**). Despite the short duration of TGF-β stimulation, the TGF-β pathway itself was also significantly upregulated. We found that *JUNB* was one of the most upregulated genes (**Fig. 3d, e and g**). In contrast, in the HTLV-1–infected non-leukemic ATL7 cell line, TGF-β stimulation has almost no effect on gene expression level, and GSEA did not identify any significantly enriched pathways (**Fig. 3f, g**). This indicates that the ATL7 cell line is insensitive to TGF-β stimulation. We next examined the expression of *JUNB* at the mRNA level in various ATL cell lines with or without TGF-β treatment. After two hours of stimulation with TGF-β, *JUNB* expression was increased in both ATL cell lines (ED and MT-1). In contrast, in a non-leukemic HTLV-1–infected T-cell line (ATL7), TGF-β stimulation slightly decreased *JUNB* expression (**Fig. 3h**). To further understand the mechanism by which TGF-β stimulation increases *JUNB* expression, we performed ChIP-seq analysis for pSmad3 after TGF-β activation. In the ATL cell lines ED and MT-1, pSmad3 bound specifically to the *JUNB* promoter region, suggesting that pSmad3 directly promotes *JUNB* transcription. Thus ATL cell lines respond to TGF-β in the expected manner: pSmad3 enters the nucleus and induces *JUNB* transcription. Although pSmad3 has the ability to bind to the *JUNB* promoter region in non-leukemic ATL7 cells, it does not appear to induce *JUNB* expression (Supplementary Fig. 4).

**Fig. 3.**
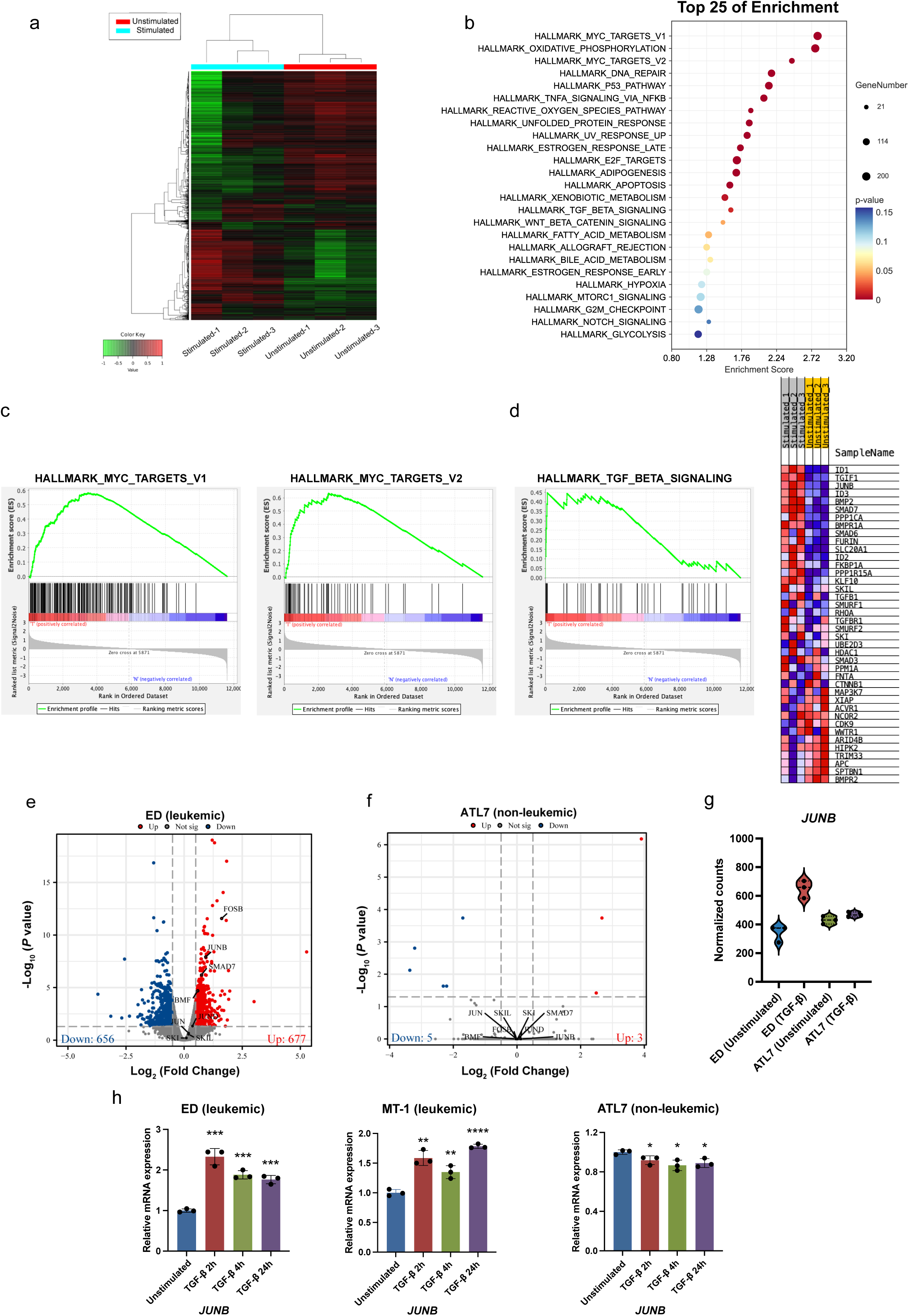
TGF-β-dependent transcriptional regulation in the ED cell line. ED cells were subjected to RNA-seq with or without 2-hour TGF-β treatment. **a** Heatmap of relative mRNA expression levels in TGF-β stimulated ED cells vs. unstimulated ED cells. **b** Hallmark gene sets upregulated in ED cells upon TGF-β treatment, analyzed by Gene Set Enrichment Analysis (GSEA). **c** Myc-associated gene signatures are significantly enriched in TGF-β stimulated ED cells compared with unstimulated ED cells, as determined by GSEA with RNA-seq. **d** Significantly enriched gene signatures for the TGF-β related gene set (left) and enriched genes (right) were determined by GSEA. **e** Volcano plot representing genes differentially expressed in TGF-β stimulated ED cells vs. unstimulated cells. Genes considered significantly upregulated or downregulated are highlighted in red and blue, respectively. Genes with insignificant changes are highlighted in grey. **f** Volcano plot representing genes differentially expressed in TGF-β treated ATL7 cells vs. unstimulated cells. **g** Normalized counts of *JUNB* in TGF-β treated ED cells and ATL7 cells vs. control cells. **h** *JUNB* mRNA levels in ED, MT-1 and ATL7 cell lines without or with TGF-β treatment for 2, 4 or 24 hours were analyzed by RT-qPCR (triplicate experiments, * p < 0.05, ** p < 0.01, *** p < 0.001, **** p < 0.0001).

Recent studies have shown that like Smad3, JunB can shuttle between the cytoplasm and the nucleus^24^ and is a known binding partner of HBZ protein^25^. These facts suggest that JunB may also play a role in the altered HBZ localization after TGF-β stimulation. To validate this hypothesis, we first verified the binding of JunB and HBZ protein by co-immunoprecipitation experiments (Supplementary Fig. 3a) and Duolink^®^ PLA. In the ATL cell line ED, the HBZ-JunB complex was predominantly expressed in the nucleus. After stimulation with TGF-β, the nuclear localization of these complexes increased further (**Fig. 4a, upper panel, Fig. 4b**). In the non-leukemic HTLV-1– infected T-cell line ATL7, JunB-HBZ binding was also found mainly in the nucleus, but we did not observe any significant change in localization after TGF-β treatment (**Fig. 4a, lower panel, Fig. 4b**). Furthermore, in two clinical samples from acute ATL cases, translocation of JunB-HBZ complexes was also observed (**Fig. 4c, d**).

**Fig. 4.**
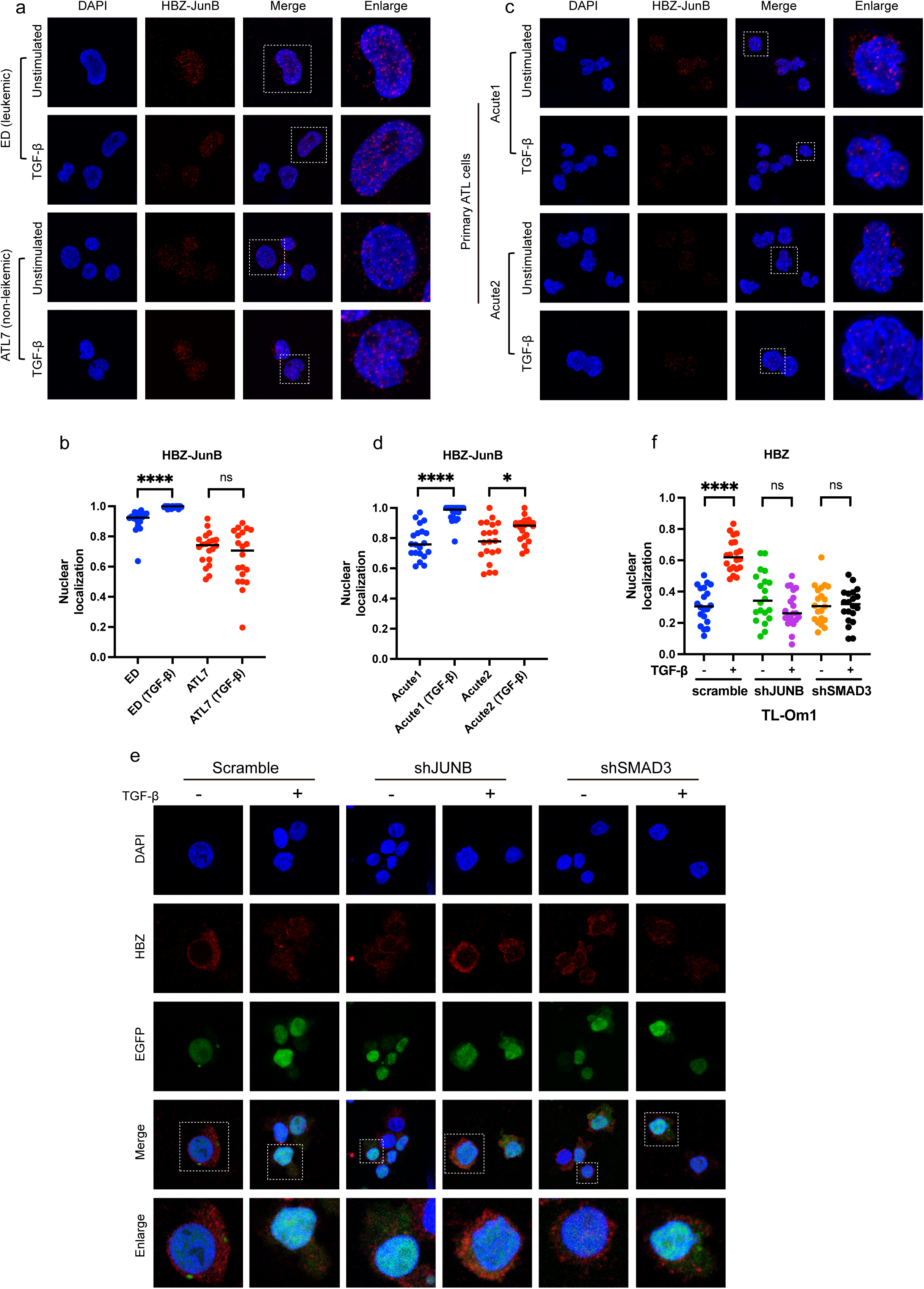
JunB plays a role in the translocation of HBZ to the nucleus induced by TGF-β treatment. **a** Complexes of JunB with HBZ in ED and ATL7 cell lines with or without TGF-β treatment were detected by Duolink^®^ PLA. **b** The proportion of HBZ-JunB complexes that are localized to the nucleus in TGF-β treated and untreated ED and ATL7 cell lines was analyzed by ImageJ. 20 randomly selected cells were analyzed for each cell line. **c** HBZ-JunB complexes in primary ATL cells of ATL patients with acute-type with or without TGF-β treatment were detected by Duolink^®^ PLA. **d** The proportion of HBZ-JunB complexes that are localized to the nucleus in primary ATL cells of acute-type ATL patients with or without TGF-β stimulation was analyzed by ImageJ. 20 randomly selected cells were analyzed for each sample. **e** Localization of HBZ in TL-Om1 cells infected with scramble shRNA, shJUNB or shSMAD3 lentivirus vectors, with or without TGF-β stimulation. **f** The proportion of HBZ protein that is localized to the nucleus in (**e**) was analyzed by ImageJ. 20 randomly selected cells were analyzed for each sample. Statistical analyses were performed by two-tailed unpaired Student’s t test. * p < 0.05, **** p < 0.0001, ns. p > 0.05. All experiments were performed at least twice.

We further examined whether JunB and Smad3 are essential for the nuclear translocation of HBZ. To this end, we knocked down the *JUNB* gene or the *SMAD3* gene by short hairpin RNA (shRNA) in the ATL cell lines TL-Om1 and ED to observe whether TGF-β stimulation could still induce translocation of HBZ protein. Compared to the scramble shRNA–infected control group in which HBZ translocated to the nucleus upon TGF-β stimulation as observed previously, the knockdown of either *JUNB* or *SMAD3* prevented HBZ from moving to the nucleus in response to TGF-β stimulation (**Fig. 4e**, **f**, and Supplementary Fig. 5a, b).

Taken together, these data demonstrate that the TGF-β-induced nuclear translocation of HBZ depends on JunB and Smad3 forming complexes with HBZ and thus facilitating its entry into the nucleus.

### *JUNB* knockdown promotes apoptosis in ATL cell lines

To further understand the function of JunB in ATL pathogenesis, we transduced lentivirus vectors expressing shRNA against *JUNB* into ED cells, MT-1 cells, and the two non-leukemic HTLV-1–infected cell lines ATL6 and ATL7 (Supplementary Fig. 6a, b). We performed a cell proliferation assay to observe the effect of *JUNB* suppression on ATL cell proliferation. In the ED and MT-1 cell lines, *JUNB* inhibition significantly inhibited ATL cell proliferation (**Fig. 5a**). In contrast, inhibition of *JUNB* in the two non-leukemic clone cell lines did not affect their cell growth, suggesting that proliferation of these non-leukemic cell lines was not dependent on *JUNB* (**Fig. 5b**).

**Fig. 5.**
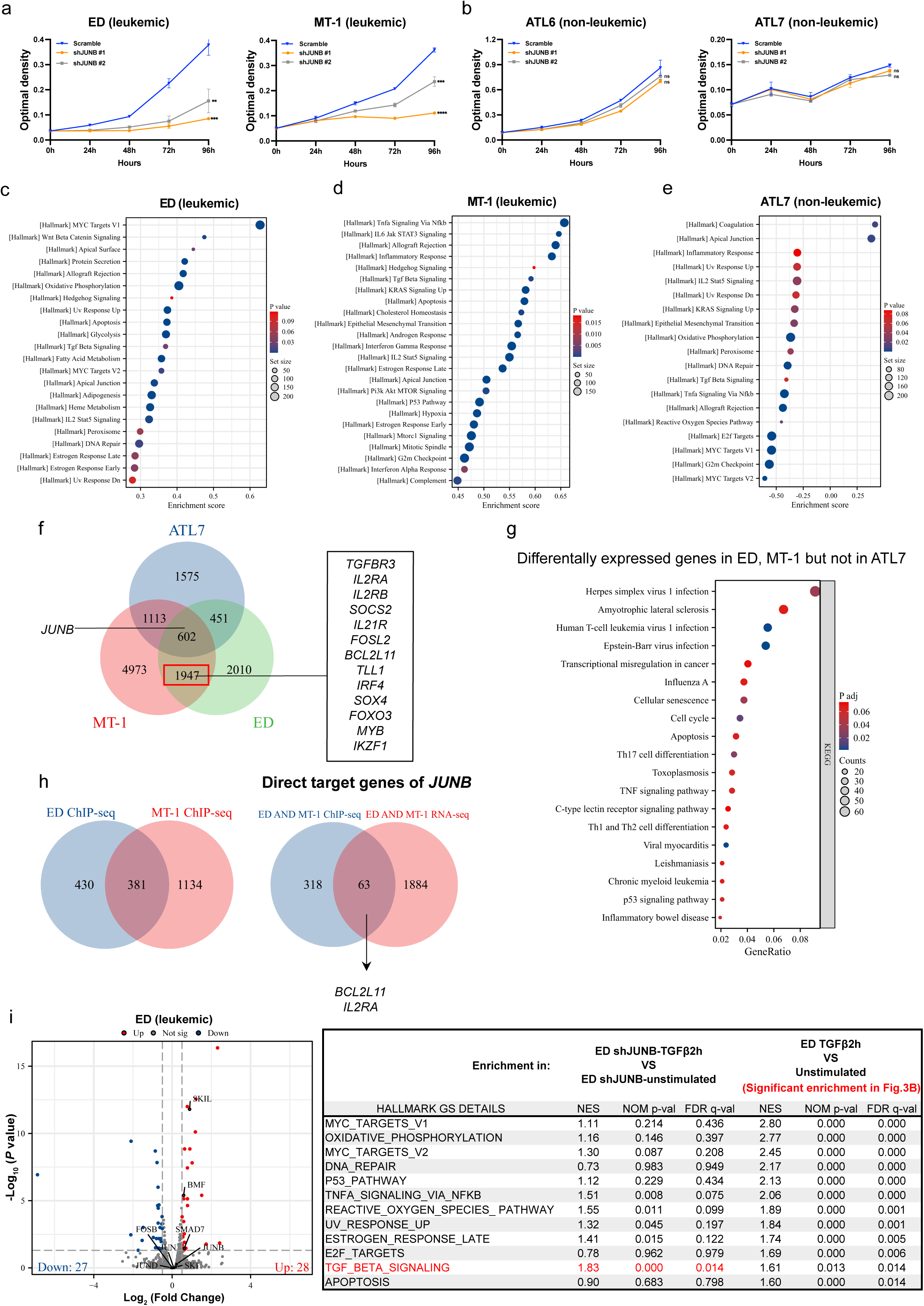
JunB-dependent transcriptional regulation in ATL cells. **a** Cell growth of ED (left) and MT-1 (right) cells infected with shJUNB#1 or shJUNB#2 (normalized mean ± s.d. of triplicate experiments). **b** Cell growth of ATL6 (left) and ATL7 (right) cells infected with shJUNB#1 or shJUNB#2 (normalized mean ± s.d. of triplicate experiments). **c-e** Significantly enriched gene signatures in *JUNB* knockdown cells, determined by Gene Set Enrichment Analysis (GSEA) with RNA-seq in ED (**c**), MT-1 (**d**) and ATL7 (**e**) cells. **f** Venn diagram showing the overlap of differentially expressed genes identified in *JUNB* knockdown cells vs, control cells in ED, MT-1 and ATL7 cells. **g** Kyoto Encyclopedia of Genes and Genomes (KEGG) analysis of genes that are found to be differentially expressed (*JUNB* knockdown cells vs. control cells) in ED and MT-1 but not ATL7 cells. **h** Venn diagram showing the overlap of JunB target genes (by ChIP-seq) and genes found by RNA-seq to be differentially expressed in both ED and MT-1 cells, but not in ATL7 cells. **i** Volcano plot (left) showing genes differentially expressed in TGF-β stimulated *JUNB* knockdown ED cells compared with unstimulated knockdown cells. Genes considered significantly upregulated or downregulated are highlighted in red and blue, respectively. Genes with insignificant changes are highlighted in grey. The table (right) shows the GSEA of gene expression after TGF-β stimulation for *JUNB* knockdown cells and parental ED cells. Gene sets are significantly enriched at FDR < 0.05.

To further understand the mechanism by which *JUNB* suppression inhibited the growth of ATL cell lines, we performed RNA-seq in two ATL cell lines (ED and MT-1) and a non-leukemic HTLV-1-infected cell line (ATL7), each transduced with shJUNB or scramble shRNA. Notably, the apoptosis related gene set was significantly upregulated in both the ED and MT-1 cell lines (**Fig. 5c**, **d**, and Supplementary Fig. 6c). In contrast, the apoptosis-related gene set was not significantly altered in the ATL7 cell line (**Fig. 5e**). Moreover, we identified 2054 genes whose expression was significantly changed in the ED and MT-1 cell lines, but not in the ATL7 cell line (**Fig. 5f**). KEGG analysis also revealed that these genes were mainly associated with the cell cycle and apoptosis (**Fig. 5g**). These results were consistent with the results of the proliferation experiments, where suppression of *JUNB* expression affected the ATL cell lines but not the non-leukemic cells. Thus *JUNB* has leukemic cell-specific functions critical for the persistence of ATL cells.

To elucidate the potential mechanisms of JunB-dependent transcriptional regulation in ATL cell lines vs. the HTLV-1–infected non-leukemic cell line, we performed ChIP-seq analysis for JunB in ED, MT-1 and ATL7 cells infected with shJUNB or scramble shRNA. To focus on genes directly regulated by JunB, we looked for genes that had JunB ChIP-seq peaks and were differentially expressed (upregulated or downregulated) by knockdown of *JUNB* (**Fig. 5h**). Thus we defined a set of 63 genes directly regulated by JunB specifically in both ED and MT-1 cells. *BCL2L11*, which is a pro-apoptotic BCL family gene, was one of the most enriched genes in the apoptosis gene set in the ED and MT-1 cell lines in our RNA-seq data. *JUNB* knockdown induced expression of *BCL2L11*, suggesting that this upregulation led to apoptosis in ATL cells (Supplementary Fig. 7, left). Additionally, *JUNB* regulated a set of regulatory T cell-related genes, such as *IL2RA*, *IL2RB*, and *SOCS2*, in ATL cell lines (**Fig. 5f**). Moreover, *JUNB* directly promoted the expression of *IL2RA* in ATL cell lines (Supplementary Fig. 7, right). In contrast, in the non-leukemic cell line, the expression level of *BCL2L11* is not regulated by JunB, which might explain why the proliferation of non-leukemic cells is not affected by the loss of *JUNB*. Thus the development of ATL appears to involve JunB becoming a regulator of genes that it previously did not regulate in HTLV-1 carrier cells.

We also evaluated the role of *JUNB* in the TGF-β signaling pathway of ATL cells. We knocked down *JUNB* in ED cells and performed RNA-seq before and after TGF-β stimulation (**Fig. 5i, left**). Compared to parental ED cells (**Fig. 3e**), the number of differentially expressed genes under TGF-β stimulation was significantly reduced when *JUNB* was knocked down, and those genes that were up-regulated, such as *SKIL*, *SMAD7*, and *BMF*, were known negative regulators of the TGF-β/Smad signaling pathway^26, 27^. Furthermore, GSEA showed that only the TGF-β signaling pathway was significantly upregulated when *JUNB* was knocked down. In contrast, pathways enriched by TGF-β stimulation in parental ED cells (**Fig. 3b**), such as cell growth-related signaling pathways and the MYC pathway, were not significantly enriched when *JUNB* was knocked down (**Fig. 5i, right**). Similar results were also observed in the MT-1 cell line (Supplementary Fig. 8a, b). These results demonstrate that *JUNB* is an important activator of the TGF-β signaling pathway in ATL cells.

Collectively, through a combined analysis of RNA-seq and ChIP-seq, we defined the set of genes directly regulated by JunB in ATL cells and demonstrated that JunB is an important activator of the TGF-β signaling pathway in ATL cells. We also found that JunB activates gene sets associated with cell proliferation and inhibits the expression of the pro-apoptotic gene *BCL2L11*. Importantly, only in ATL cells, but not in non-leukemic infected cells, does JunB serve as a core regulator. Thus JunB plays a distinct and critical role in the oncogenesis of ATL.

### *JUNB* is critical for ATL cell growth *in vivo*

A previous study has shown that knockdown of *HBZ* gene expression significantly reduces the size and infiltration of ATL tumors *in vivo*^10^. Given the core regulatory role of the JunB-HBZ axis in ATL oncogenesis, we were interested in testing the regulatory function of JunB in an *in vivo* environment. To this end, we knocked down the *JUNB* gene in two ATL cell lines, MT-1 and ED, and injected these cells subcutaneously into NOD/SCID/γ-chain knockout (NSG) mice to observe the growth of tumor cells (**Fig. 6a**). In both cell lines, all five mice in the control groups without *JUNB* knockdown successfully developed tumors. However, in the shJUNB-MT-1 injected group, we did not observe tumor formation in any of the five mice (**Fig. 6b-d**). In the shJUNB-ED injected group, *JUNB* knockdown significantly inhibited tumor size and tumor growth compared to the control mice (**Fig. 6e-g**). These results suggest that JunB plays an indispensable role in ATL tumorigenesis *in vivo*.

**Fig. 6.**
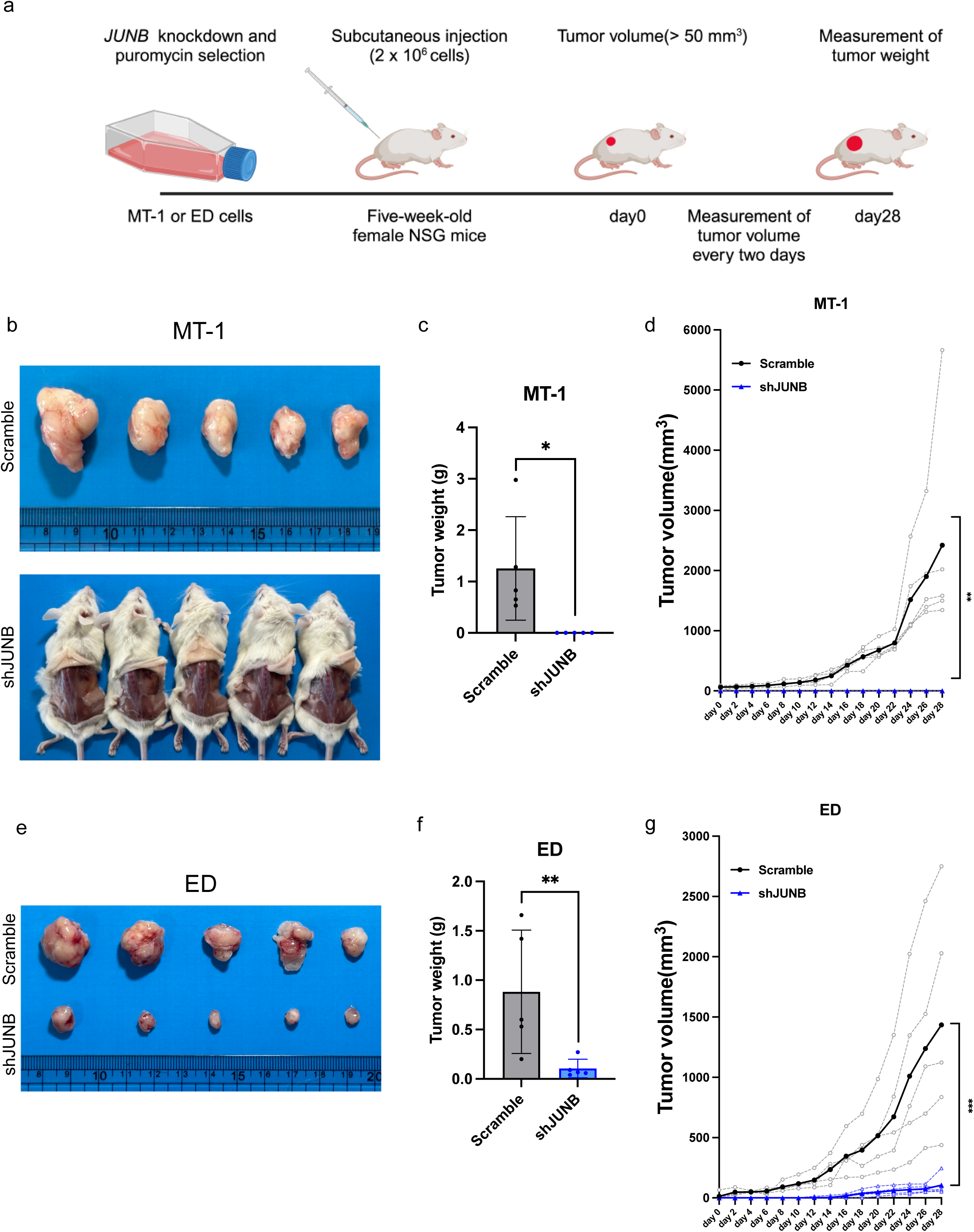
JunB depletion leads to ATL tumor growth inhibition *in vivo*. **a** Schematic representation of the xenograft model of ATL. **b** ATL tumor growth in mice subcutaneously injected with scramble shRNA (n = 5) or shJUNB (n = 5) infected MT-1 cells. **c, d** ATL tumor weights (normalized mean ± s.d) (**c**) and volumes (**d**) in the mice subcutaneously injected with scramble shRNA (n = 5) or shJUNB (n = 5) infected MT-1 cells. **e** ATL tumor growth in mice subcutaneously injected with scramble shRNA (n = 5) or shJUNB (n = 5) infected ED cells. **f, g** ATL tumor weights (normalized mean ± s.d) (**f**) and volumes (**g**) in the mice subcutaneously injected with scramble shRNA (n = 5) or shJUNB (n = 5) infected ED cells. Statistical analysis is by two-tailed unpaired Student’s t test; * p < 0.05, ** p < 0.01, *** p < 0.001).

## Discussion

*HBZ*, which is a central viral gene in HTLV-1 pathogenesis, is consistently expressed not only in ATL cells but also in infected cells from HTLV-1 carriers; however, the functional differences of HBZ in these different contexts remain unclear. Elucidating the mechanisms underlying these differences is crucial for understanding ATL oncogenesis. In this study, we found that nuclear localization of HBZ protein is higher in ATL cells than in non-leukemic HTLV-1–infected cells. Furthermore, TGF-β activation causes HBZ protein translocation from the cytoplasm into the nucleus only in ATL cells but not in non-leukemic infected cells, suggesting that TGF-β-induced nuclear localization of HBZ is associated with ATL oncogenesis. It remains unclear why HTLV-1–infected cells that are not leukemic fail to respond to TGF-β in a similar manner. Discovering the “trigger” that converts an infected cell from TGF-β nonresponsive to responsive is an important issue that needs to be clarified in the future.

It is known that HBZ is involved in the regulation of multiple oncogenic pathways by interacting with many transcription factors and transcription co-activators, such as Smad proteins, CBP/p300^12, 13^, and the AP-1 proteins^25, 28, 29^. Additionally, HBZ also regulates the activity of transcription factors. For example, HBZ drives the expression of BATF3 in ATL cells, regulating the BATF3/IRF4 transcriptional network^30^. These findings support the crucial functional role of nuclear HBZ in ATL oncogenesis. In particular, co-localization of JunB with HBZ in the nucleus is thought to be critical for ATL development, as shown in this study.

Previous research has reported that JunB is expressed in all ATL cases and that the loss of *JUNB* is toxic to ATL cell lines^31^. However, the mechanism of JunB regulation in ATL remains unclear. We demonstrate here that TGF-β upregulates the expression of *JUNB* and induces JunB-HBZ complex translocation to the nucleus in ATL cells but not in non-leukemic cell lines. Additionally, the cytotoxicity resulting from the knockdown of *JUNB* is only present in ATL cell lines, while non-leukemic cell lines continue to proliferate normally in the absence of *JUNB*, indicating that *JUNB* is a master transcription factor for oncogenic properties of ATL cells. JunB has been reported to play a crucial role in the proliferation of helper T cell subsets by inhibiting the expression of the pro-apoptotic gene *BCL2L11* (encoding Bim)^32^. In this study, we found that JunB suppresses Bim expression by binding directly to the *BCL2L11* genomic region, thereby protecting ATL cells from apoptosis. However, in the non-leukemic HTLV-1–infected cell line, this regulatory function of JunB was absent and *JUNB* deficiency did not have cytotoxic effects.

ATL cells have immunophenotypes that closely resemble that of iTreg cells, due to the induction of the *FOXP3* gene by HBZ^33^. It is well known that naïve T cells differentiate into iTregs by inducing Foxp3 through TGF-β stimulation^34^. Previously, we reported that HBZ-Tg mice have an increased number of iTregs with high Foxp3 expression and develop inflammation and T-lymphomas^11^. Those findings suggest that expansion of Tregs triggered by HBZ accelerates oncogenesis in this subset. Notably, JunB has been reported to promote the expression of Treg effector molecules^35^ and Treg differentiation through the IL-2 pathway^36^. Our RNA-seq and ChIP-seq data suggest that JunB directly upregulates *IL2RA* expression to increase sensitivity to IL-2, thereby promoting the formation of the Treg phenotype in ATL cells. Previous studies showed that like JunB, HBZ targets these genes, *BCL2L11*^37^ and *IL2RA*^30^. Indeed, it has been shown that HBZ suppresses transcription of *BCL2L11* in the nucleus^37^. It is noteworthy that super-enhancers are present at the *IL2RA* locus in ATL cells^30^, suggesting that the JunB and HBZ proteins are recruited to the locus and coordinately regulate the transcription of multiple genes during ATL oncogenesis.

Interestingly, recent study has shown that, in breast cancer cells, *JUNB* is a late-responding gene induced by TGF-β stimulation and creates a feed-forward regulatory network that enhances their invasive capacity^38, 39^. Here we show that activation of the TGF-β signaling pathway causes pSmad3 to be recruited to the *JUNB* promoter region, where it directly stimulates *JUNB* expression in ATL cells, but not in non-leukemic HTLV-1–infected cells. Meanwhile, pSmad3 and JunB also facilitate the nuclear translocation of HBZ, where it exerts its oncogenic functions; our findings suggest that HBZ expands leukemic cells by reinforcing the link between TGF-β signaling and *JUNB*-induced cell proliferation, which, as we have shown, becomes critical for the survival of ATL cells.

Besides HTLV-1, other human oncogenic retroviruses also encode proteins that promote infection and oncogenic processes through various mechanisms in the nucleus. In human papillomavirus (HPV), the E6 and E7 proteins interfere with cell cycle regulation by affecting nuclear proteins such as the tumor protein p53 (TP53) and retinoblastoma (Rb)^40^. In Epstein–Barr virus (EBV), Epstein–Barr nuclear antigen 1 (EBNA-1) helps maintain viral DNA in the nucleus and regulates viral gene expression, while the latent membrane protein 1 (LMP1) acts in the nucleus to promote cell proliferation^41^. In addition, the nuclear localization of the hepatitis B virus (HBV)-encoded oncogene X protein (HBx) is essential for promoting HBV replication, thereby contributing to the development of hepatocellular carcinoma (HCC)^42, 43, 44^. The subcellular localization of HBx may be influenced by its abundance ^45^ and interactions with cellular proteins^46, 47^. However, there is no clear consensus on the preferential subcellular localization of these viral proteins in viral oncogenesis. Our results confirm that viral proteins may achieve nuclear translocation under signaling pathway stimulation with the help of binding proteins. Identification of suitable binding proteins could provide insight into the subcellular translocation mechanisms of other viral proteins.

In conclusion, our study demonstrates that the nuclear localization of HBZ protein is associated with disease status, and is likely influenced by differences in responsiveness to TGF-β. Upon TGF-β stimulation, cytoplasmic HBZ can bind with pSmad3 or JunB and translocate into the nucleus in ATL cells (Supplemental Fig. 9). Having entered the nucleus, HBZ can further act to upregulate the TGF-β pathway, creating a sort of positive feedback loop that maintains this pathway in an activated state. In addition, HBZ and JunB act to upregulate other genes associated with proliferation and the Treg phenotype. The result would be leukemic cells that are constantly driven to proliferate — a crucial step in oncogenesis. This discovery of the Smad3-JunB-HBZ cascade induced by TGF-β could pave the way for development of therapeutic strategies against this fatal malignancy.

## Methods

### Clinical samples

All patients were fully informed about the purpose and procedures of this study. Written informed consent was obtained in accordance with the Declaration of Helsinki and received approval from the Institutional Review Board of Kumamoto University (Genome 297). Peripheral blood mononuclear cells (PBMCs) were isolated from patients with adult T-cell leukemia-lymphoma (ATL) and human T-cell leukemia virus type 1 (HTLV-1) carriers by Ficoll-Paque PLUS (GE Healthcare).

### Cell lines

Human embryonic kidney cell lines HEK293T purchased from American Type Culture Collection were cultured in Dulbecco’s modified Eagle’s medium (Nacalai Tesque) supplemented with 10% fetal bovine serum (FBS) and antibiotics. HTLV-1 immortalized cell lines (MT-2, MT-4 and Hut102) and ATL cell lines (MT-1, ED, TL-Om1, ATL43T[+]) were cultured in RPMI 1640 (Nacalai Tesque) with 10% FBS and antibiotics. HTLV-1–infected non-leukemia cell lines (ATL6, ATL7) and an ATL cell line (ATL55T [+]) were cultured in RPMI 1640 (Nacalai Tesque) with 10% FBS, recombinant human IL-2 and antibiotics.^8^

### Mice

NOD.Cg-PrkdcscidIl2rgtm1Wjl /SzJ (NSG) mice were purchased from Charles River Laboratories Japan, Inc. All animal experiments were performed in accordance with Kumamoto University Animal Care and Use Committee guidelines and were approved by the Committee.

### Duolink® Proximity Ligation Assays

Cell lines or PBMCs extracted from patient blood samples were incubated with or without recombinant human TGF-β1 (Wako, 20ng/ml) for 1 hour at 37°C. Cells were washed twice with Dulbecco’s Phosphate-Buffered Saline (D-PBS) and adjusted to the appropriate cell number, then concentrated onto glass coverslips (Matsunami Glass) by Cytospin (Shandon). Cells were then fixed in 4% paraformaldehyde for 15 minutes at room temperature, permeabilized with 0.2% Triton X-100 for 15 minutes at room temperature, and blocked with Duolink® Blocking Solution for 30 minutes in 37°C, then stained with mouse anti-HBZ antibody (1A10, generated by immunizing C57BL/6 mice with keyhole limpet hemocyanin [KLH]-conjugated HBZ peptide 97-133 [CKQIAEYLKRKEEEKARRRRRAEKKAADVARRKQEEQE]) and either rabbit anti-HBZ polyclonal antibody, generated by immunizing with HBZ-peptides (CRGPPGEKAPPRGETH and QERRERKWRQGAEKC) (Medical & Biological Laboratories) or JUNB Rabbit mAb (3753, Cell Signaling) or Phospho-SMAD3 (C25A9) Rabbit mAb (Cell Signaling) diluted in Duolink® Antibody Diluent (1x) overnight at 4°C. Then, cells were incubated with Duolink® In Situ PLA® Probe Anti-Mouse PLUS and Duolink® In Situ PLA® Probe Anti-Rabbit MINUS in 37 °C for 1 hour, then incubated with Duolink® 5x Ligation Buffer and Duolink® Polymerase (10U/µl) at 37°C for 30 minutes, and finally incubated with Duolink® Amplification Red (5x) and Duolink Ligase (1 U/µl) at 37 °C for 100 minutes. The glass coverslips were sealed with Duolink ® In Situ Mounting Medium with DAPI. The cells were observed with a C2 confocal microscope (Nikon) and analyzed with ImageJ. Data was collected from 20 randomly selected HBZ-expressing cells for each sample.

### Co-immunoprecipitation experiments

For endogenous immunoprecipitation experiments, MT-2 and ATL55T (+) cells were lysed in RIPA lysis buffer with protease inhibitor for 30 min on ice. Rabbit anti-HBZ polyclonal antibody (Medical & Biological Laboratories) was used for immunoprecipitation. Normal rabbit IgG (Wako) was used as a control. The lysates were immunoprecipitated with antibody conjugated Dynabeads™ Protein G Magnetic Beads (Invitrogen).

### Extraction of cytoplasmic and nuclear proteins

TL-Om1 cells were treated with or without recombinant human TGF-β1 (Wako, 7178bhf), *SMAD3*(VB900050-5901tdd, VB900050-5907vsa) and the negative control were purchased from Vector Builder. Lentiviral vectors were generated by transient transfection of HEK293T cells. The lentiviral vectors were harvested 48 hours after transfection and used immediately for infection. After 24 hours of infection, target cells were transferred into fresh growth medium, and the stably transfected cells were selected in puromycin.

### RNA extraction and Quantitative RT-PCR

Total RNA was extracted using the ReliaPrep RNA Miniprep System (Promega) following the manufacturer’s instructions. 1 μg of total RNA was synthesized into cDNA with Superscript IV Reverse Transcriptase (Thermo Fisher Scientific). Semiquantitative RT-PCR was performed using FastStart Universal SYBR Green Master (Roche) and the StepOnePlus Real-Time PCR System (Applied Biosystems). The primers used were as follows: JUNB-forward 5’-CGATCTGCACAAGATGAACCACG-3’, JUNB-reverse 5’-CTGCTGAGGTTGGTGTAAACGG-3’.

### Cell proliferation assay

After knockdown of *JUNB* and selection in puromycin, cells were plated at an appropriate cell density in 96-well plates. The Cell Counting Kit-8 (DOJINDO LABORATORIES) was used according to the manufacturer’s instructions to assess the proliferation of cells.

### RNA-sequencing

For RNA sequencing, each experiment contained 3 biological replicates. RNA samples were extracted using the ReliaPrep RNA Miniprep System (Promega). Total RNA was provided for library preparation and sequencing to Macrogen, which used the TruSeq Stranded mRNA Library Prep Kit and NovaSeq6000 (Illumina). After cleaning the row reads with Trimmomatic^48^, reads were mapped to hg38 by STAR. Differential gene expression was analyzed by RSEM^49^ and edgeR^50^. Differentially expressed genes (log2 Fold change < -0.5 or > 0.5, FDR< 0.05) were further analyzed using Gene Set Enrichment Analysis by GSEA software^51^ or KEGG (Kyoto Encyclopedia of Genes and Genomes) enrichment analysis^52^.

### ChIP-sequencing

After cells were treated with or without TGF-β for 2 hours, cells were collected and ChIP DNAs were prepared using the SimpleChIP Enzymatic Chromatin IP Kit (Cell Signaling Technology) according to the manufacturer’s instructions. For Chromatin IP, JunB Rabbit mAb (Cell Signaling Technology) or Phospho-SMAD3 (C25A9) Rabbit mAb (Cell Signaling Technology) was used. DNA sequencing libraries were generated on the Illumina NovaSeq6000 platform using the TruSeq ChIP Library Preparation Kit (Illumina) with a standard 150-bp paired-end read protocol. Fastq files were trimmed and qualified with Trimmomatic and Fastqc. Trimmed reads were mapped using Bowtie2 aligned to the human genome (hg19)^53^. Reads that aligned twice or more and unassembled reads (chrM, random and chrUn) were removed by using samtools^54^. PCR duplicates were also deleted with Picard (MarkDuplicates). Subsequently, by using samtools with the “-q 30” option, we selected reads with a mapping quality of more than 30. The ENCODE Blacklist genome regions were also deleted by using BEDtools^55^. ChIP-seq peaks were assessed by MACS2^56^ with the “-p 1e-5” option using the corresponding input-sequenced file as a control. Extracted peaks were then annotated to the nearest genes of the hg19 by using ChIPseeker^57^. To visualize deep-sequencing data, we used samtools, deeptools^58^ and SparK (https://github.com/harbourlab/SparK).

### Xenograft experiments

ED and MT-1 cell lines were infected with Scramble shRNA or shJUNB lentivirus, followed by puromycin selection. Two million cells for each cell line were injected subcutaneously into 6-week-old female NSG mice. When the tumor size became approximately 50 mm³, the subcutaneous tumor burden was measured every two days using calipers. Tumor volumes (mm^3^) were calculated using the following formula: 0.5 x (longest length) x (width)^2^. On day 14 after the beginning of measurement, the mice were sacrificed, and tumor weight was measured. In general, animals were sacrificed when their tumors reached 2cm or when the mice became moribund.

### Statistical analysis and data visualization

All data was statistically analyzed using the two-tailed unpaired Student’s t test using Prism 10 (GraphPad Software). All experiments were performed with at least three biological replicates. The minimum significance level was set at P < 0.05. Asterisks indicate the statistical significance as follows: *P < 0.05; **P < 0.01; ***P < 0.001; **** P < 0.0001. Schematic diagrams in this study were created with BioRender.com.

### Data and materials availability

Data of deep sequence, RNA-seq, and ChIP-seq in this study will be deposited at the DDBJ database and be publicly available as of the date of publication.

## Acknowledgements

We thank Miho Matsumoto for technical support, Daisuke Kurotaki for technical advice on bioinformatic analyses, and Linda Kingsbury for proofreading. This study was supported by grants from the Project for Cancer Research And Therapeutic Evolution (P-CREATE) (20cm0106306h0005 to M.M., J.Y.); the Research Program on Emerging and Re-emerging Infectious Diseases of the Japan Agency for Medical Research and Development (AMED), (20fk0108088h0002 and 24fk0108629h0003 to M.M. and J.Y.); the Science and Technology Platform Program for Advanced Biological Medicine, also of AMED (21am0401003h0003 to J.Y.); the Japan Society for the Promotion of Science (JSPS) KAKENHI (19H03689 to M.M., 23H02936 to J.Y. 23K15303 to T.S.); the JSPS Core-to-Core Program A, Advanced Research Networks to M.M., J.Y.; the Center for Metabolic Regulation of Healthy Aging (CMHA) to T.S.; and JST SPRING, Grant Number JPMJSP2127 to W.Z..

## Author Contributions

W.Z. acquired funding for the project, performed almost all of the experiments, (including Duolink PLA, gene expression profile, functional analysis, and cell culture assays), performed bioinformatic analysis and data curation, and wrote the paper. T.S. conceived and supervised the project, acquired funding for the project, performed gene expression profile, data curation and bioinformatic analysis, provided clinical samples, and wrote the paper. X.C. provided assistance with the xenograft experiments. M.M. supervised the project, acquired funding for the project, performed data curation. J.Y. conceived and supervised the project, acquired funding for the project, performed data curation, provided clinical samples, and wrote the paper. All authors discussed the results and commented on the manuscript.

