## Supporting Information for "JunB-HBZ nuclear translocation by TGF-β is a key driver in HTLV-1–mediated leukemogenesis"

**This PDF file includes:**
Figures S1 to S9

a

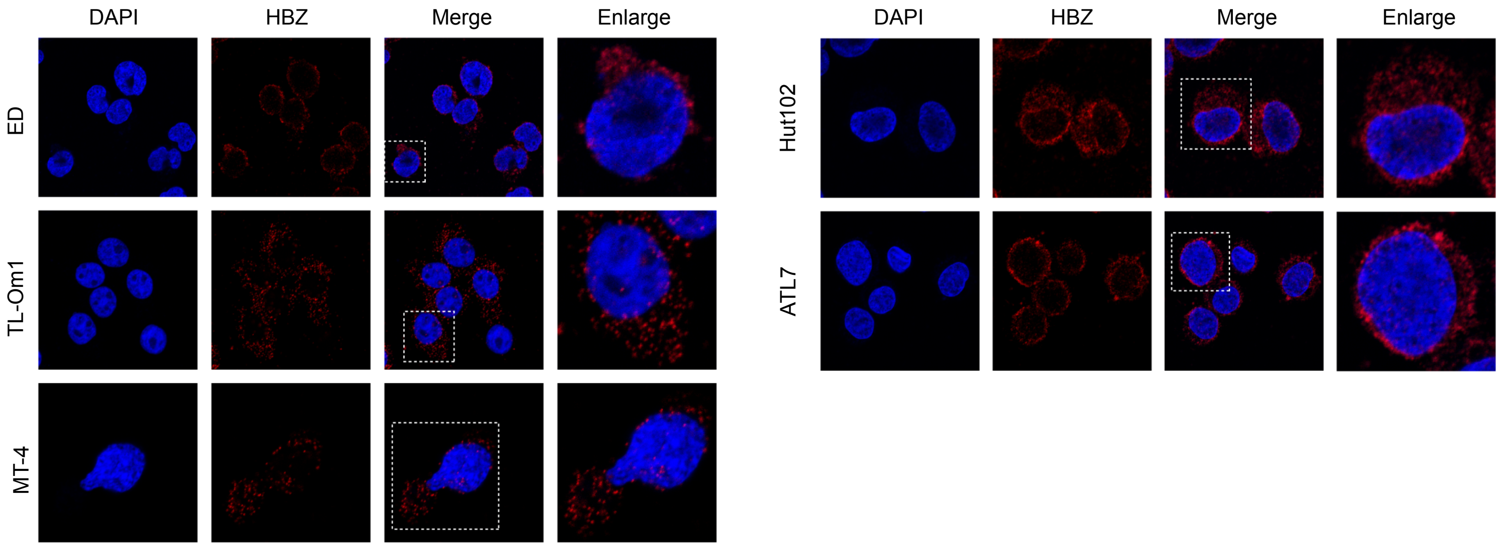


b

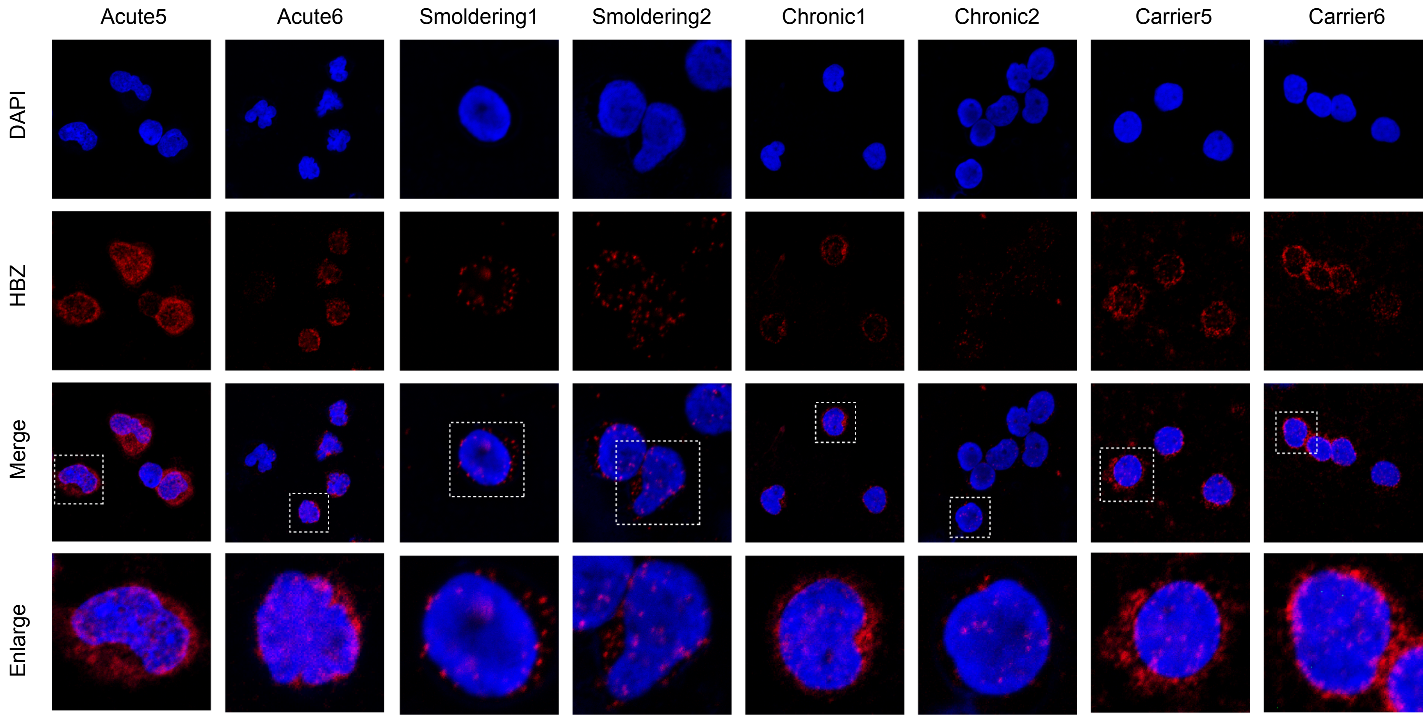


c

Characteristics of patients

| Patient | Proviral load* | Nuclear localization of HBZ |
| --- | --- | --- |
| acute1 | 115.9 | 0.659 |
| acute2 | 14.2 | 0.569 |
| acute3 | 177.9 | 0.241 |
| acute4 | 101.2 | 0.515 |
| acute5 | 96.6 | 0.575 |
| acute6 | 94.6 | 0.600 |
| smoldering1 | 7.8 | 0.764 |
| smoldering2 | 12.1 | 0.623 |
| chronic1 | 12.8 | 0.655 |
| chronic2 | 92.9 | 0.815 |
| carrier1 | 19.8 | 0.451 |
| carrier2 | 5.9 | 0.419 |
| carrier3 | 12.6 | 0.301 |
| carrier4 | 32.7 | 0.184 |
| carrier5 | 8.1 | 0.296 |
| carrier6 | 11.8 | 0.384 |

*Proviral load (copies per 100 PBMCs) is measured by real-time PCR.

**Supplementary Figure 1. Expression pattern of HBZ protein in various cell lines and patient samples.**

**a** Localization of HBZ protein in ED, TL-Om1, MT-4, Hut102 and ATL7 cell lines by Duolink® PLA. **b** Localization of HBZ protein in ATL cells from ATL patients with acute-type, smoldering-type and chronic-type, and HTLV-1–infected cells of HTLV-1–infected carrier patient samples. **c** Provirus load and the proportion of HBZ protein that is localized to the nucleus were analyzed in ATL patients with acute-type, smoldering-type and chronic-type and HTLV-1–infected carrier patients. All experiments were performed at least twice.

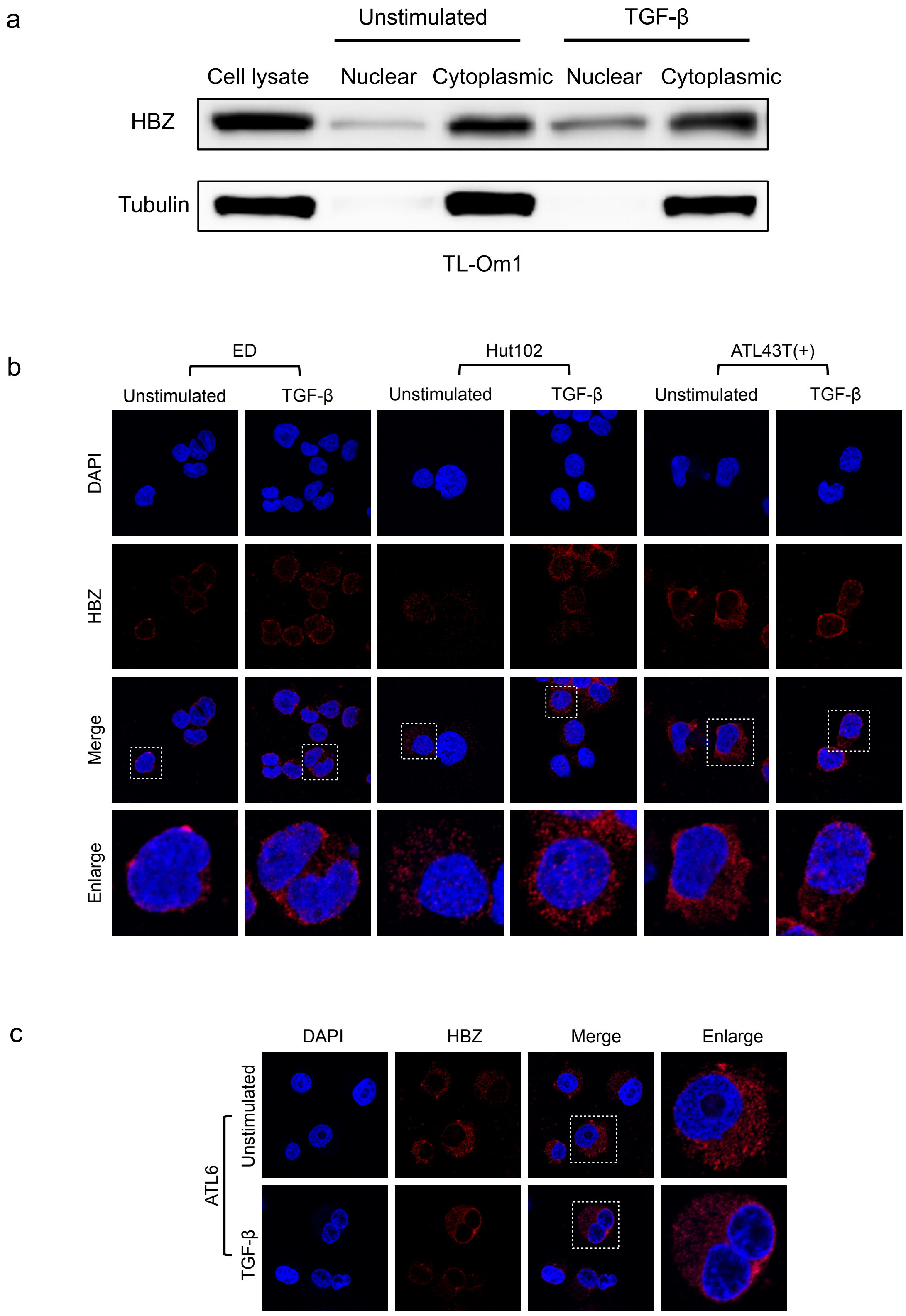


**Supplementary Figure 2. HBZ translocates into the nucleus after TGF-β stimulation in ATL cells.**

**a** Immunoblotting of HBZ in cytoplasmic and nuclear fractions in TL-Om1 cells. Tubulin is a control for cytoplasmic fractions. **b** Localization of HBZ protein in ED, Hut102 and ATL43T (+) cell lines with or without TGF-β stimulation was detected by Duolink® PLA. **c** Localization of HBZ protein in the HTLV-1–infected non-leukemic clone cell line ATL6 with or without TGF-β stimulation. All experiments were performed at least twice.

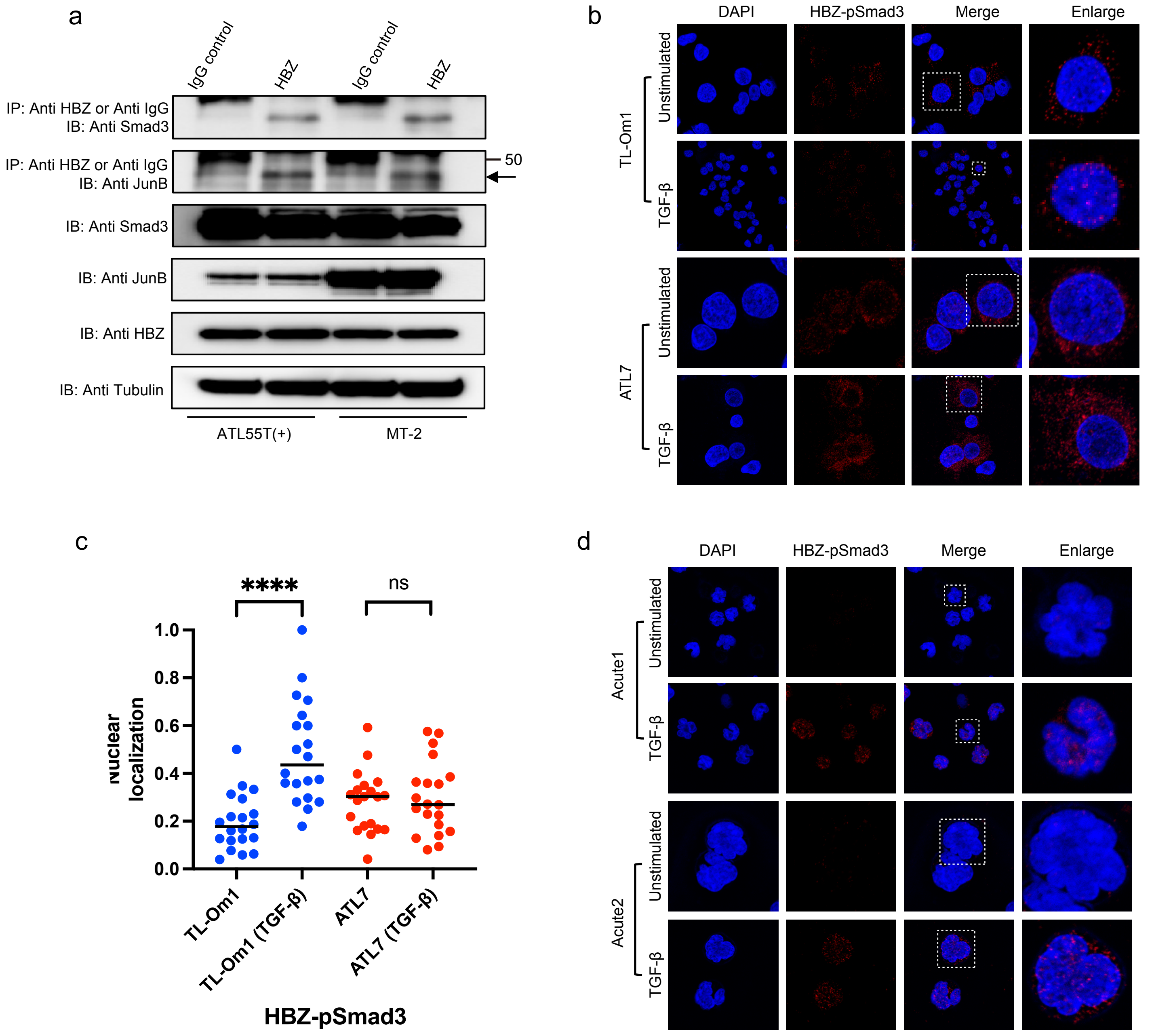


**Supplementary Figure 3. TGF-β treatment induces translocation of HBZ-pSmad3 complexes in ATL.**

**a** Co-immunoprecipitation experiment showing the Smad3-HBZ interaction and the JunB-HBZ interaction in ATL55T (+) and MT-2 cell lines. IP, immunoprecipitation; IB, immunoblot. **b** Complexes of phosphorylated Smad3 (pSmad3) with HBZ in TL-Om1 and ATL7 cell lines with or without TGF-β treatment were detected by Duolink® PLA. **c** The proportion of HBZ-pSmad3 complexes that are localized to the nucleus in TL-Om1 and ATL7 cell lines with or without TGF-β treatment was analyzed by ImageJ. 20 randomly selected cells were analyzed for each cell line. **d** HBZ-pSmad3 complexes in primary ATL cells from acute-type ATL patients, with or without TGF-β stimulation. Statistical analyses were performed by two-tailed unpaired Student's t test. **** $p < 0.0001$, ns. $P > 0.05$. All experiments were performed at least twice.

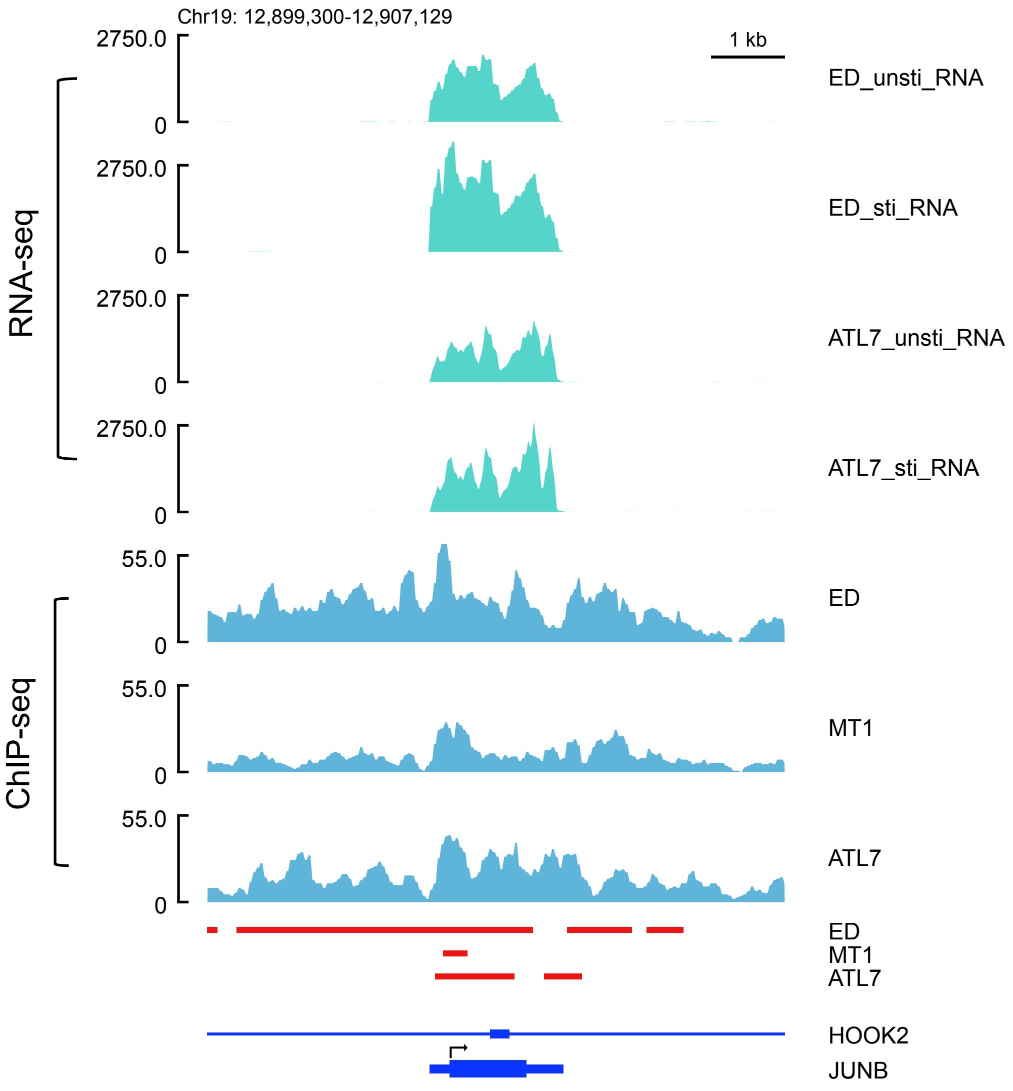


**Supplementary Figure 4. pSmad3 directly regulates the *JUNB* gene in ATL cell lines.** RNA-seq analysis of ED and ATL7 cells stimulated with TGF-β for two hours compared with their unstimulated counterparts, and pSmad3 enrichments (ChIP-seq) of the *JUNB* gene in ED, MT-1 and ATL7 cell lines. Peaks of pSmad3 ChIP-seq are shown as red bars in the *JUNB* gene.

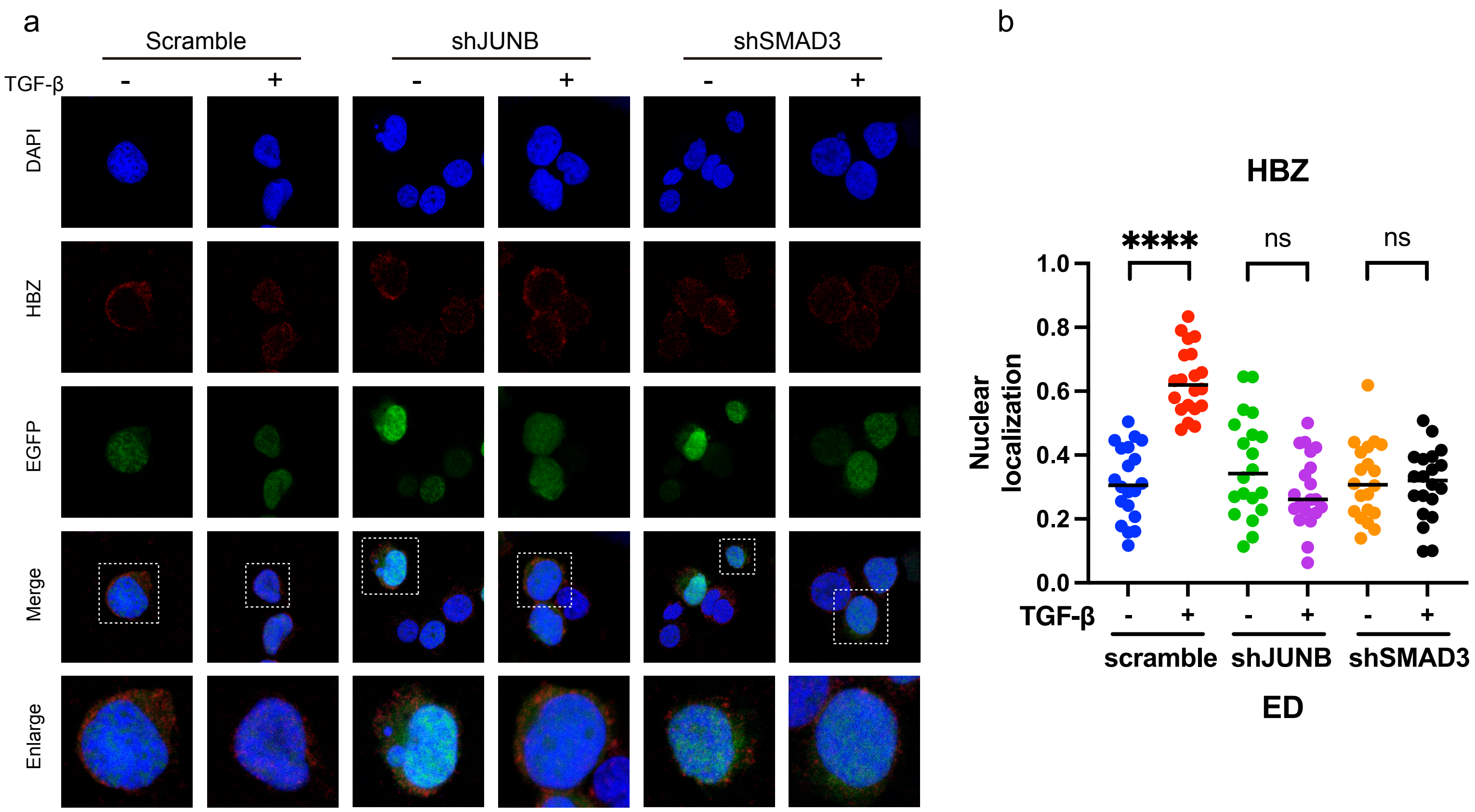


**Supplementary Figure 5. pSmad3 and JunB are required for the TGF-β-induced nuclear translocation of HBZ in ED cells.**

**a** Localization of HBZ in ED cells transduced with lentivirus vectors expressing shRNA against scramble, *JUNB* or *SMAD3* with or without TGF-β stimulation, detected by Duolink® PLA. **b** The proportion of HBZ that is localized to the nucleus was analyzed by ImageJ. 20 randomly selected cells were analyzed for each cell line. Statistical analyses were performed by two-tailed unpaired Student's t test. **** $p < 0.0001$, ns. $P>0.05$. All experiments were performed at least twice.

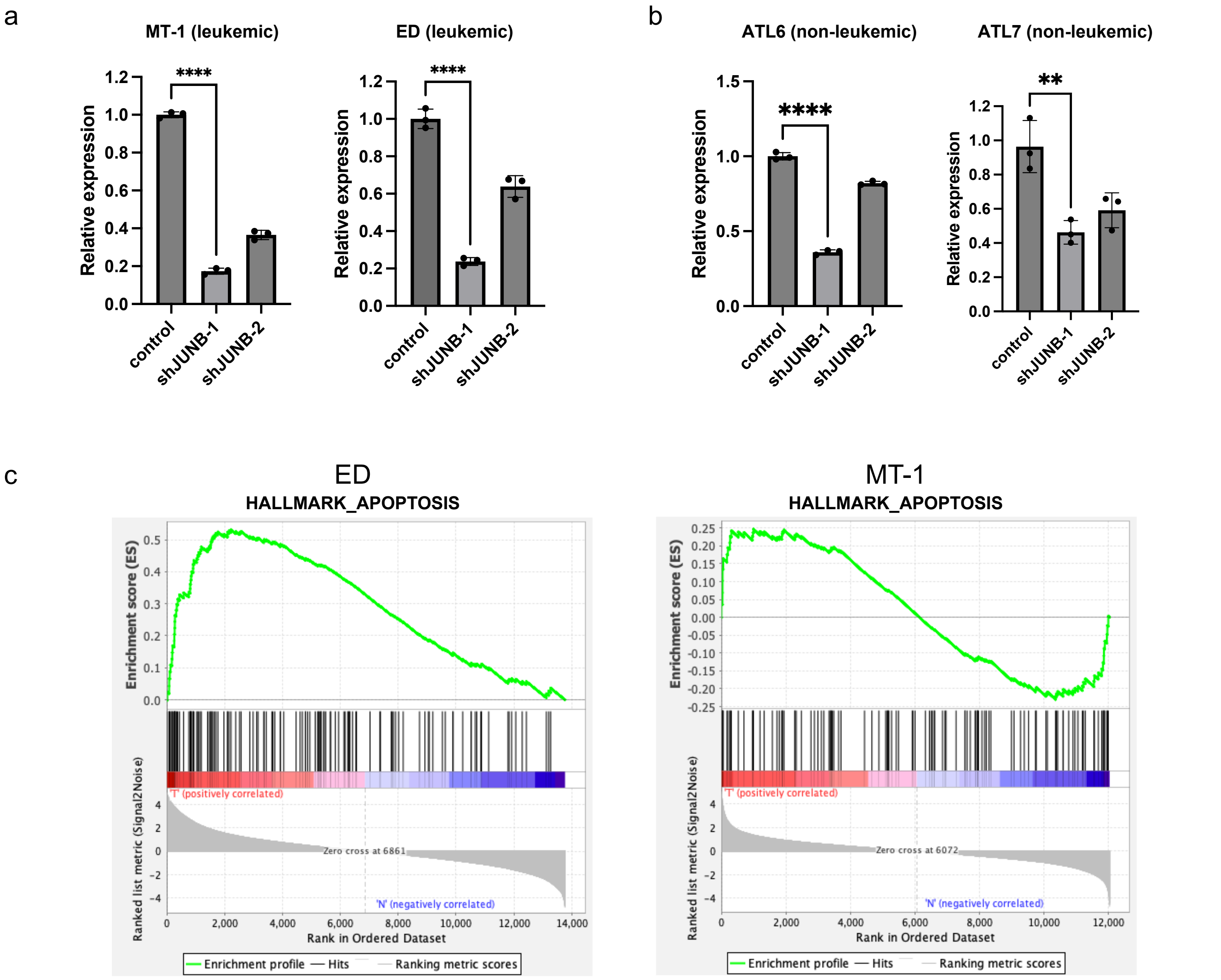


**Supplementary Figure 6. Apoptosis gene sets were upregulated in *JUNB* knockdown ATL cells.**

**a-b** *JUNB* mRNA levels in the ED and MT-1 cell lines (**a**) and in the ATL6 and ATL7 cell lines (**b**) transduced with scramble shRNA, shJUNB-1 or shJUNB-2 were analyzed by RT-qPCR after puromycin selection (triplicate experiments, Statistical analyses were performed by two-tailed unpaired Student's t test. ** $p < 0.01$, **** $p < 0.0001$). **c** Significantly enriched gene signatures for apoptosis in *JUNB* knocked down cells, determined by GSEA analysis with RNA-seq in ED and MT-1 cells.

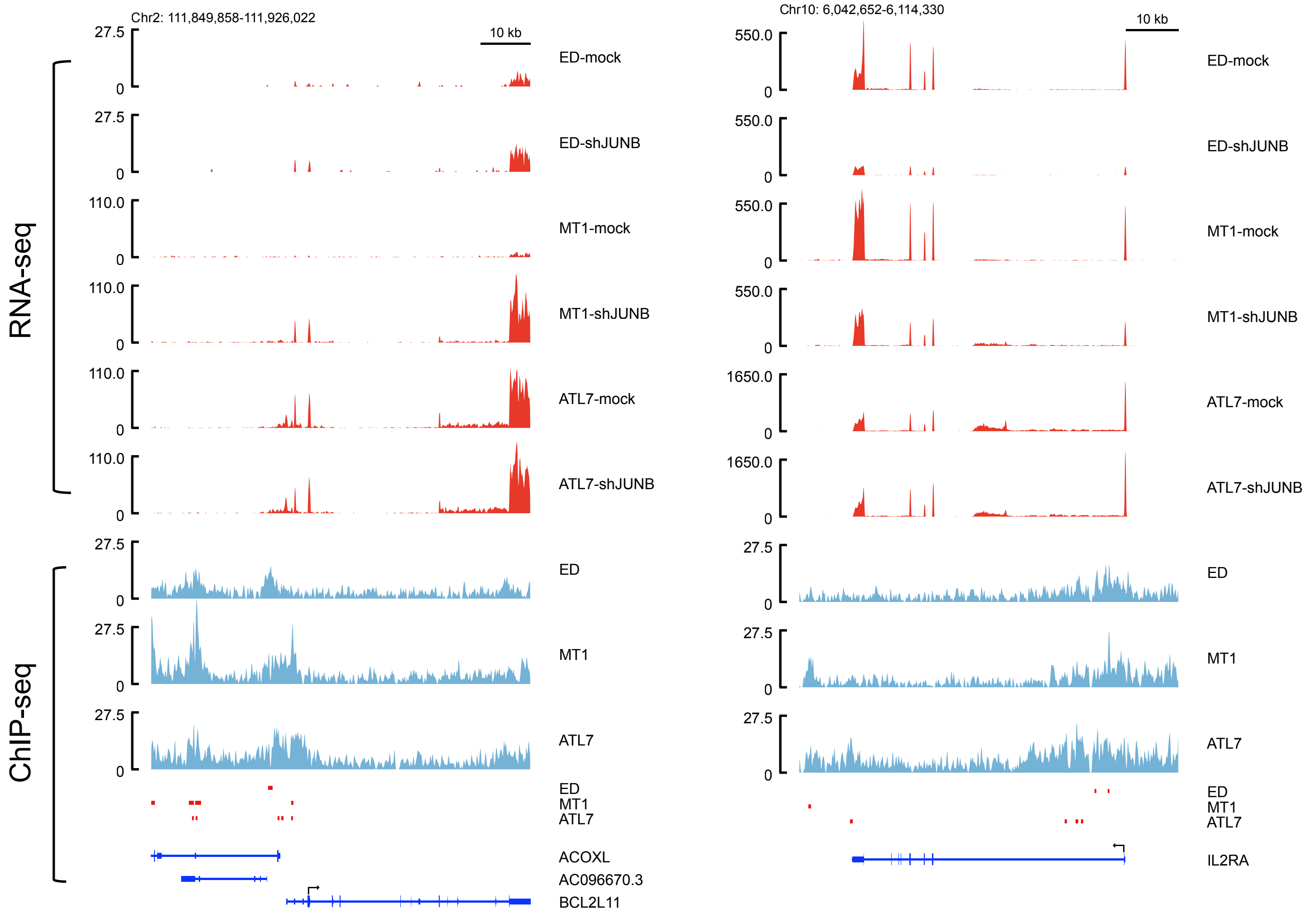


**Supplementary Figure 7. JunB directly regulates the *BCL2L11* and *IL2RA* genes in ATL cells.** JunB enrichment (ChIP-seq) and transcripts (RNA-seq) of the *BCL2L11* (left) and *IL2RA* (right) genes in ED, MT-1 and ATL7 cells transduced with scramble shRNA or shJUNB. Peaks of JunB ChIP-seq are shown as red bars in both genes.

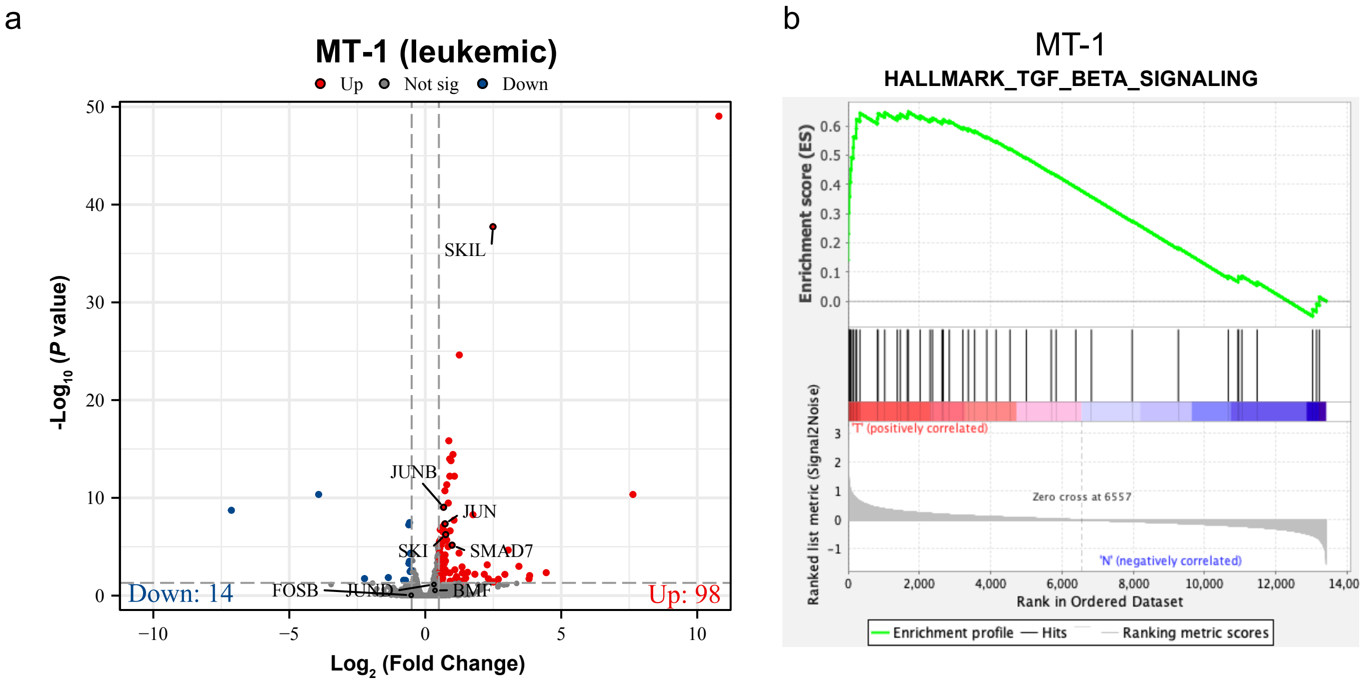


**Supplementary Figure 8.** ***JUNB*** **is an important activator of the TGF-β signaling pathway in ATL cells.**

**a** Volcano plot representing genes differentially expressed in TGF-β stimulated *JUNB* knockdown MT-1 cells compared with unstimulated knockdown cells. Genes that are significantly upregulated or downregulated by TGF-β stimulation are highlighted in red and blue, respectively. Genes with insignificant changes are highlighted in grey. **b** Significantly enriched gene signatures for TGF-β gene sets in *JUNB* knocked down cells, determined by GSEA analysis with RNA-seq in MT-1 cells.

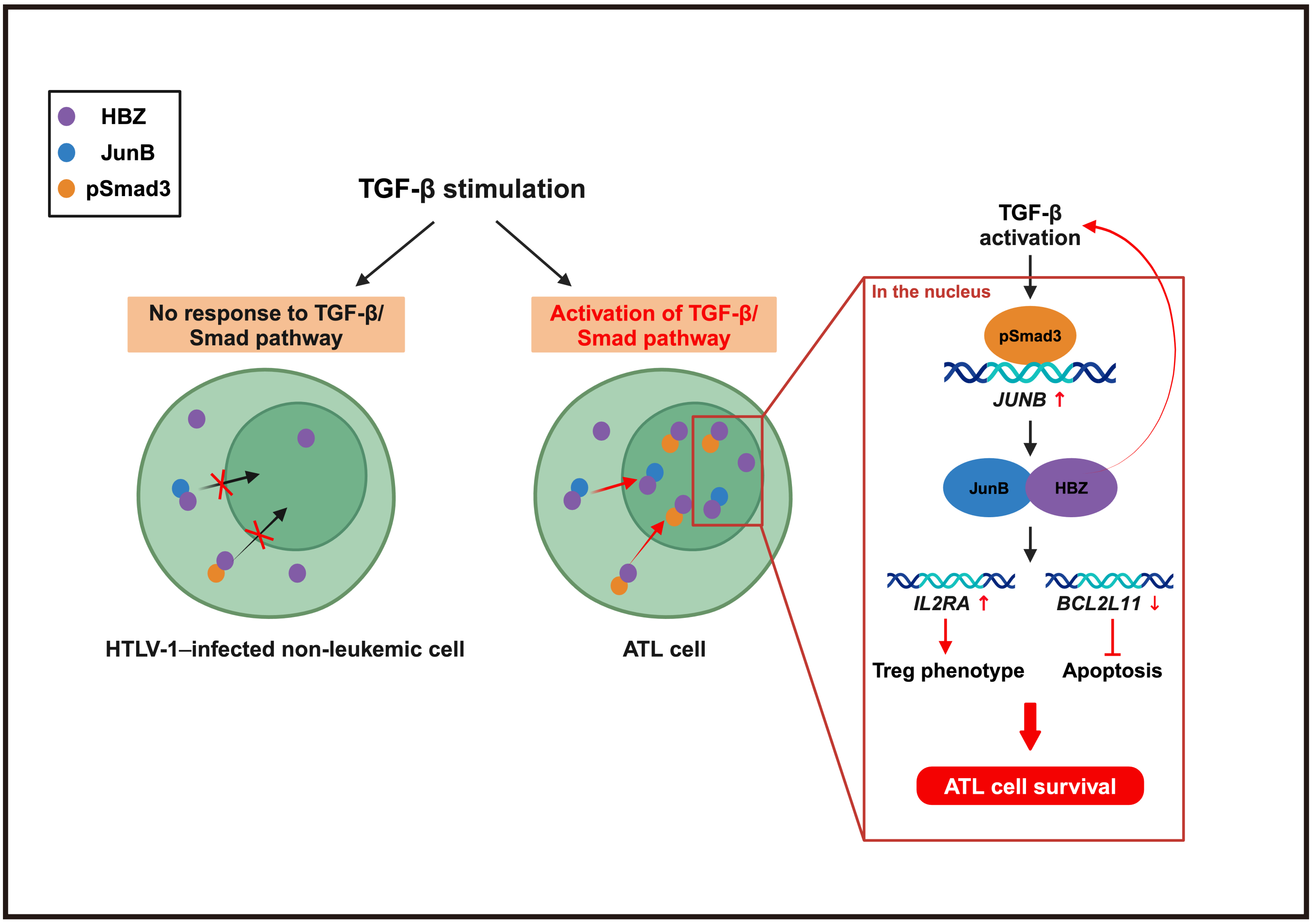


**Supplementary Figure 9. Schema of the differences in the Smad3-JunB-HBZ axis under TGF-β stimulation in ATL cells vs. HTLV-1–infected non-leukemic cells.**

HBZ is present at a higher ratio in the nucleus in ATL cells than in HTLV-1–infected non-leukemic cells. Under TGF-β stimulation, JunB and pSmad3 facilitate the nuclear translocation of HBZ in ATL cells. pSmad3 directly promotes *JUNB* expression, and HBZ further enhances TGF-β activation. The JunB-HBZ complex coordinately regulates the expression of multiple genes, promoting the survival of ATL cells. In contrast, non-leukemic HTLV-1–infected cells show no response to TGF-β activation, JunB and pSmad3 fail to facilitate the nuclear translocation of HBZ under TGF-β stimulation.
